# Genetic load and extinction in peripheral populations: the roles of migration, drift and demographic stochasticity

**DOI:** 10.1101/2021.08.05.455207

**Authors:** Himani Sachdeva, Oluwafunmilola Olusanya, Nick Barton

## Abstract

We analyse how migration from a large mainland influences genetic load and population numbers on an island, in a scenario where fitness-affecting variants are unconditionally deleterious, and where numbers decline with increasing load. Our analysis shows that migration can have qualitatively different effects, depending on the total mutation target and fitness effects of deleterious variants. In particular, we find that populations exhibit a genetic Allee effect across a wide range of parameter combinations, when variants are partially recessive, cycling between low-load (large-population) and high-load (sink) states. Increased migration reduces load in the sink state (by increasing heterozygosity) but further inflates load in the large-population state (by hindering purging). We identify various critical parameter thresholds at which one or other stable state collapses, and discuss how these thresholds are influenced by the genetic vs. demographic effects of migration. Our analysis is based on a ‘semi-deterministic’ analysis, which accounts for genetic drift but neglects demographic stochasticity. We also compare against simulations which account for both demographic stochasticity and drift. Our results clarify the importance of gene flow as a key determinant of extinction risk in peripheral populations, even in the absence of ecological gradients.

## Introduction

Most outcrossing populations carry a substantial masked mutation load due to recessive variants, which can contribute significantly to inbreeding depression in peripheral isolates or after a bottleneck. The extent to which the increased segregation (or fixation) of deleterious mutations due to drift (“drift load”) exacerbates extinction risk in isolated populations has been a subject of long-standing interest [1, 2, 3, 4]. Theory predicts that moderately deleterious mutations contribute the most to genetic load and extinction in small populations [2, 5]; however, the prevalence of such deleterious variants of mild or moderate effect and their dominance values remain poorly characterised, except for a few model organisms [6, 7].

The relative risks posed by mutation accumulation and demographic stochasticity to a population depend crucially on its size, with theory suggesting that these may be comparable for populations in their thousands [1]. Additionally, environmental stochasticity — catastrophic events, as well as fluctuations in growth rates and carrying capacities, may dramatically lower extinction times [8]. Both demographic and environmental fluctuations, in turn, reduce the effective size of a population, making it more prone to fix deleterious alleles; the consequent reduction in fitness further depresses size, pushing populations into an ‘extinction vortex’, which is often characterised by a complex interaction between the effects of genetic drift, demographic stochasticity and environmental fluctuations [9].

Peripheral populations at the edges of species’ ranges receive dispersal from the core to an extent which varies over space and time. Moreover, ranges may be fragmented due to habitat loss and individual sub-populations connected to each other via low and possibly declining levels of migration. Under what conditions are such extinction vortices arrested by migration, and what are the genetic and demographic underpinnings of this effect, when it occurs?

Migration boosts numbers, mitigating extinction risk due to demographic and environmental stochasticity, or at the very least, allows populations to regenerate after chance extinction. The demographic consequences of migration are especially important in fragmented populations with many small patches [10]: above a critical level of migration, the population may survive as a whole over long timescales even if individual patches frequently go extinct [11, 12].

Migration also influences extinction risk by shifting the frequencies of fitness-affecting variants: the resultant changes in fitness may decrease or increase population size, thus further boosting or depressing the relative contribution of migration to allele frequency changes within a population, setting in motion a positive feedback which may culminate in extinction (when gene flow is largely maladaptive; e.g., [13, 14]) or evolutionary rescue (if gene flow supplies variation necessary for local adaptation, or reduces inbreeding load; e.g., [5, 15, 16]).

The maladaptive consequences of migration have largely been explored for extended populations under spatially varying environments: here, gene flow typically hinders local adaptation, especially at range limits, leading to ‘swamping’ and extinction [17]. However, the consequences of gene flow for fitness, and consequently survival, are not always intuitive when the fitness effects of genetic variants are uniformly deleterious (or beneficial) across populations. For example, while gene flow may alleviate inbreeding load by preventing the fixation of deleterious alleles in small populations, it may also render selection against recessive mutations less effective by increasing heterozygosity. A striking consequence is that under a range of conditions, the fitness of metapopulations is maximised at intermediate levels of migration [18] and more generally, at intermediate levels of population structure [19].

A key consideration is whether or not gene flow is symmetric, i.e., whether some sub-populations are merely influenced by the inflow of genes from the rest of the habitat or if all sub-populations influence the genetic composition of the population as a whole [20]. Asymmetric dispersal is common at the geographic peripheries of species’ ranges or on islands. Moreover, populations occupying small patches within a larger metapopulation with a wide distribution of patch sizes, or sub-populations with lower-than-average fitness (and consequently, atypically low numbers) may also experience predominantly asymmetric inflow of genes. Asymmetric gene flow allows for allele frequency differences across the range of a population even in the absence of environmental heterogeneity, e.g., when population sizes (and hence the efficacy of selection relative to drift) vary across the habitat. This, in turn, may generate heterosis or outbreeding depression across *multiple* loci, when individuals from different regions hybridize.

From a conceptual viewpoint, the consequences of asymmetric gene flow are typically simpler to analyse as we can focus on a single population, while taking the state of the rest of the larger habitat as ‘fixed’. Such analyses are key to understanding more general scenarios where genotype frequencies and population sizes across different regions co-evolve.

Here, we analyse the eco-evolutionary dynamics of a single island subject to migration from a larger mainland in a scenario with uniform selection across the two populations, i.e., where fitness is affected by a large number of variants that are unconditionally deleterious. We ask: under what conditions can migration from the mainland alleviate inbreeding load, thus preventing ‘mutational meltdown’ and extinction of the island population? Further, how are the effects of migration mediated by the genetic architecture of load, i.e., by the genome-wide mutation target and fitness effects of deleterious variants? A key focus is to understand the coupled evolution of allele frequencies (across multiple loci) and population size: to this end, we consider an explicit model of population growth with logistic regulation, where growth is reduced by an amount equal to the genetic load.

While the effects of maladaptive gene flow on marginal populations have been studied under various models [21, 22], there has been little work (under genetically realistic assumptions) on the (possibly) beneficial effects of migration on inbreeding load and survival. In particular, modeling the polygenic nature of fitness variation is crucial, as changes in load (e.g., due to migration) at any locus can affect all other loci by effecting changes in population size, which in turn influences the efficacy of selection across the genome.

## Model and Methods

Consider a peripheral island population subject to one-way migration from a large mainland. Individuals are diploid and carry *L* biallelic loci that undergo bidirectional mutation between the wildtype and deleterious state at rate *u* per generation per individual per locus in either direction. Mutations have the same fitness effects on the mainland and island, i.e., there is no environment-dependent fitness component.

Deleterious variants across loci affect fitness multiplicatively (no epistasis): individual fitness is 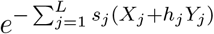, where *X*_*j*_(*Y*_*j*_) equals 1 if the *j*^*th*^ locus is homozygous (heterozygous) for the deleterious allele, and is zero otherwise. Here, *s*_*j*_ is the homozygous selective effect and *h*_*j*_ the dominance coefficient for the deleterious allele at locus *j*. We assume 0≤*h*_*j*_≤1/2, so that deleterious alleles are (partially) recessive. In individual based-simulation (see Appendix A, SI), we use the form (1−*hs*)^*n*^(1−*s*)^*m*^ (which is equivalent to the above fitness function for small *s*) where *n* represents the number of heterozygous loci and m represent the number of loci homozygous for the deleterious allele.

In each generation, a Poisson-distributed number of individuals (with mean *m*_0_) migrate from the mainland to the island. For simplicity, we assume that the mainland population is large enough that deleterious allele frequencies among migrants (denoted by 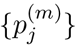) are close to the deterministic predictions for a single locus under mutation-selection equilibrium.

We assume density-independent selection on individuals on the island and density-dependent population growth with logistic regulation, where the baseline growth rate is reduced by the genetic load: the population size *n*_*t*_ in any generation *t* is then Poisson-distributed with mean equal to 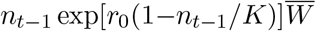, where *n*_*t*−1_ is the population size in the previous generation, *r*_0_ the baseline growth rate, *K* the carrying capacity, and 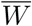 the mean genetic fitness. The *n*_*t*_ individuals in the *t*^*th*^ generation are formed by randomly sampling 2*n*_*t*_ parents (with replacement) from the *n*_*t*−1_ individuals in the previous generation with probabilities equal to their relative fitnesses, followed by free recombination between parental haplotypes to create gametes. Diploid offspring are then formed by randomly pairing gametes.

### Simulations

We carry out two kinds of simulations: individual-based simulations that explicitly track multi-locus genotypes of all individuals on the island, and simulations that assume LE (linkage equilibrium) and neglect inbreeding. The latter kind of simulations are computationally less intensive as they only track allele frequencies at the *L* loci and the size of the population. However, they make two simplifying assumptions: first, that genotypes at any locus are in Hardy-Weinberg proportions (i.e., no inbreeding); second, that any statistical associations, e.g., between the allelic states of different loci (linkage disequilibria or LD) or between the probability of identity by descent, and consequently homozygosity, at different loci (identity disequilibria or ID) — are negligible. Then, individual genotypes are simply random assortments of deleterious and wildtype alleles, and can be generated, e.g., in a simulation, by independently assigning alternative allelic states to different loci with probabilities equal to the allele frequencies. Details of the simulations are provided in Appendix A, SI.

Since selection pressures are identical on the mainland and island, systematic differences in allele frequencies or homozygosity between the two populations across multiple loci (which would generate LD and ID respectively) must arise solely due to differences in population size, which would cause the efficacy of selection to be different on the mainland and island. In general, we expect LD and ID to be negligible when all ecological and evolutionary processes are slower than recombination [14]: this may not hold, however, when populations are small (and drift significant), making it necessary to evaluate how associations between deleterious variants affect extinction thresholds. In the rest of the paper, we will only show results of the allele frequency simulations (assuming LE and zero inbreeding); we compare these with individual-based simulations and discuss how LD and ID affect population outcomes in Appendix B (SI).

### Joint evolution of allele frequencies and population size

Assuming LE and no inbreeding, the evolution of the island population is fully specified by how allele frequencies {*p*_*j*_} and the population size *n* co-evolve in time *t*. If all evolutionary and ecological processes (except recombination) are slow, we can describe this co-evolution in continuous time. We rescale all rates by the baseline growth rate *r*_0_ and population sizes by the carrying capacity *K*. This yields the following dimensionless parameters: *τ*=*r*_0_*t*, *S*=*s/r*_0_, *M*_0_=*m*_0_/(*r*_0_*K*), *U*=*u/r*_0_, *N*_*t*_=*n_t_/K* and an additional parameter *ζ*=*r*_0_*K* which governs the strength of demographic fluctuations. The joint evolution of allele frequencies {*p*_*j*_} and the scaled population size *N* in continuous time is described by the following equations (see also [14]):

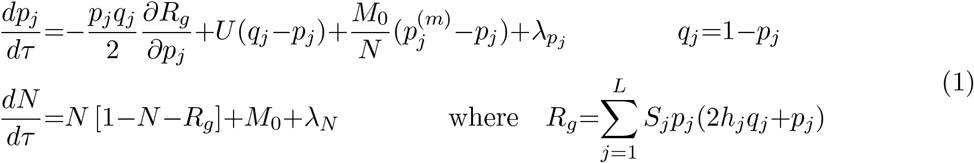

The random processes *λ*_*N*_ and 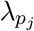 satisfy 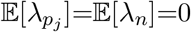, 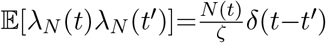 and 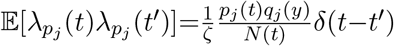, where 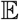 denotes the mean. The four terms in the first equation correspond (in order of appearance) to changes in allele frequency due to selection, mutation, migration and drift. Note that the strength of migration is inversely proportional to population size *N*, reflecting the stronger (relative) effect of migration on the genetic composition of smaller, as opposed to larger, island populations. The second equation describes the evolution of population size *N*: the first term describes changes in *N* under logistic growth, where the growth rate is reduced by a factor proportional to the log mean fitness (i.e., the genetic load); the second term captures the effect of migration; the third term corresponds to demographic fluctuations (whose variance is proportional to *N*, the size of the population). These equations capture a key feature of polygenic eco-evolutionary dynamics — namely, that the evolution of allele frequencies at different loci is *coupled* via their dependence on a common *N*, which in turn is influenced by the degree of maladaptation at all loci via *R*_*g*_. Thus, allele frequencies do *not* evolve independently, even if allelic states at different loci are statistically independent at any instant (under LE).

For *fixed N* and under LE and IE, the joint distribution of allele frequencies at mutation-selection-migration-drift equilibrium is a product of the single-locus distributions. This was given by Wright [23], and for a given locus *j* is (in terms of scaled parameters):

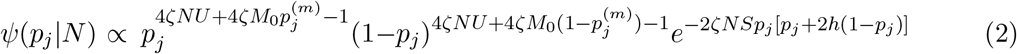

Integrating over this distribution yields the expected allele frequency 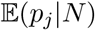 and the expected heterozygosity 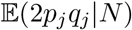 at any locus, and thence the expected total load 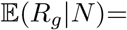 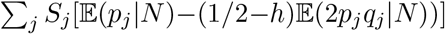 (scaled by the baseline growth rate *r*_0_) for fixed *N*.

However, in reality, *N* is not fixed and will fluctuate— both due to random fluctuations in fitness (due to the underlying stochastic fluctuations of {*p*_*j*_} at equilibrium) as well as in the reproductive output of individuals (demographic stochasticity). Thus, *N* itself follows a distribution. While we can write down an equation for the stochastic co-evolution of *N* and {*p*_*j*_}, no explicit solution for the joint equilibrium distribution is possible unless mutation rates are strictly zero (as assumed by [14]). Thus, we must employ various approximations to describe the coupled dynamics of population sizes and allele frequencies.

### Approximate semi-deterministic analysis

We expect the population size distribution to be sharply peaked around one or more values {*N*_*_} if demographic fluctuations are weak (i.e., for *ζ*=*r*_0_*K*≫1) and if fluctuations in mean fitness are also weak (i.e., for large *L*). Such sharply-peaked distributions correspond to populations that transition only rarely between alternative peaks, i.e., where the alternative peaks {*N*_*_} of the size distribution represent ‘metastable’ states, allowing allele frequencies sufficient time to equilibrate at any *N*_*_. Then, the genetic load would be close to the expected value 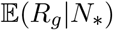 under mutation-selection-drift-migration balance (*given N*_*_) when populations are in one of the alternative metastable states, though not necessarily while they transition between states. In order to determine {*N*_*_}, we postulate that these must represent stable equilibria of the population size dynamics, neglecting demographic stochasticity and assuming that mean fitness (given *N*_*_) is close to the expectation under mutation-selection-drift-migration balance for that *N*_*_. Then we have:

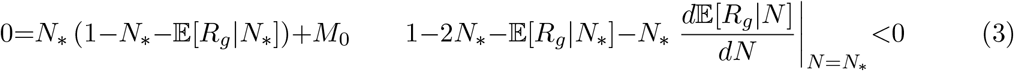

The equality follows from eq. (1), by setting the third (noise) term to zero and assuming that *R*_*g*_ is close to its *equilibrium* expectation 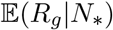, given *N*_*_. The second inequality is the condition for *N*_*_ to be a stable equilibrium: this means that populations starting at an arbitrary *N* in the vicinity of *N*_*_ would evolve towards this equilibrium size. Equation (3) can be solved numerically to obtain the equilibria {*N*_*_}, which, under the above assumptions, will be close to the peaks of the population size distribution. As we see below, depending on the parameter regime, there may be one or two stable equilibria of eq. (3), corresponding to population size distributions that are unimodal or bimodal. Bifurcations (i.e., critical parameter thresholds where one of the two stable equilibria vanishes) thus correspond to qualitative transitions in the state of the population.

We will refer to the analysis above as a ‘semi-deterministic analysis’ as it accounts for the stochastic effects of genetic drift on allele frequencies and load (via the expectation 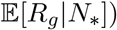) but neglects demographic stochasticity. In general, we expect the semi-deterministic analysis to become more accurate for larger *ζ*=*r*_0_*K* (see also Appendix B, SI), since increasing *ζ*, which implies higher number of births per generation, results in weaker demographic fluctuations [14].

## Results

### Metastable populations and extinction times in the absence of migration

We first consider peripheral populations in the absence of migration. Such populations necessarily become extinct in the long run: however, depending on the mutation target for deleterious mutations (relative to the baseline growth rate *r*_0_), the fitness effects of mutations and the carrying capacity of the island, extinction times may be very long and populations metastable.

We can use eq. (3) to gain intuition for the conditions for metastability under zero migration. Setting *M*_0_=0, it follows that there is always an equilibrium at *N* =0 (corresponding to extinction): this equilibrium is stable for *L*(*S/*2)>1 where *S*=*s/r*_0_, and unstable otherwise (see Appendix B for details). There may exist a second equilibrium at 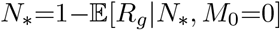; the population size *N*_*_ is positive (i.e., the population is not extinct) only if 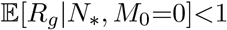, i.e., if the equilibrium genetic load is lower than the baseline growth rate.

Since the equilibrium load 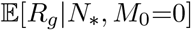 depends on four independent parameters (which we choose as *ζS*=*Ks*, *ζU* =*Ku*, *h* and 2*LU* =2*L*(*u/r*_0_)), mapping the conditions for metastability boils down to asking: in the absence of demographic fluctuations, where in this 4-dimensional parameter space, can we find populations with a non-zero equilibrium size or sufficiently low load (fig. 1(a))? In reality, there is a fifth parameter *ζ*=*r*_0_*K*, which governs demographic stochasticity: thus, all other parameters being equal, extinction times will be longer when demographic stochasticity is weaker, i.e., *r*_0_*K* larger (fig. 1(b)).

**Figure 1:**
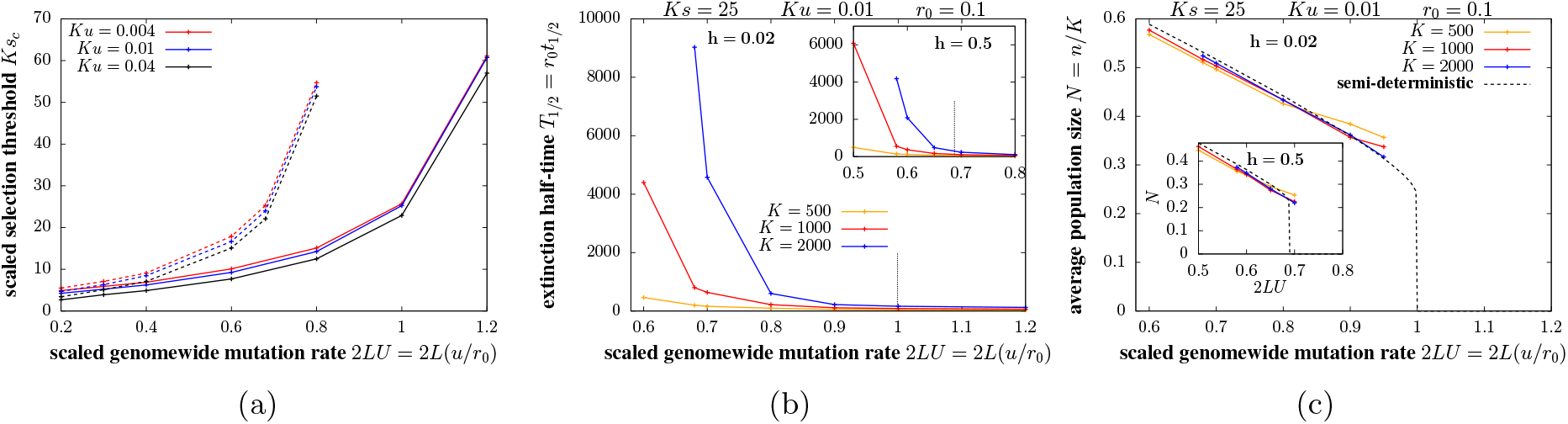
Population outcomes with zero migration. (A) Scaled critical selection threshold *Ks*_*c*_, above which populations are metastable, as a function of the scaled mutation target 2*LU* =2*L*(*u/r*_0_) for different values of *Ku* (different colors) for nearly recessive (*h*=0.02; solid lines) and additive (*h*=0.5; dashed lines) alleles. A non-zero equilibrium population size 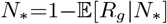 exists for *Ks>Ks*_*c*_ but not for *Ks<Ks*_*c*_. This selection threshold is calculated using eq. (3) by neglecting demographic stochasticity, and thus strictly provides a criterion for stable populations in the limit *ζ*=*r*_0_*K*→∞. (B) The scaled extinction half-time *T*_1/2_=*r*_0_*t*_1/2_ (see text for definition) as a function of 2*LU* for various *K* (different colors) for nearly recessive (*h*=0.02; main plot) and additive alleles (inset). (C) The average scaled population size *N*=*n/K* of metastable populations vs. 2*LU* for various *K* (different colors) along with the semi-deterministic prediction 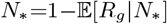 (dashed black line) for nearly recessive (*h*=0.02; main plot) and additive alleles (inset). In both (B) and (C), the carrying capacity is increased while proportionately decreasing *s*, *u* and increasing *L*, such that *Ks*=25, *Ku*=0.01 and 2*LU* =2*L*(*u/r*_0_) remain unchanged; increasing *K* thus has the sole effect of weakening demographic stochasticity. Extinction times and the average population sizes in the metastable state are computed from allele frequency simulations (under LE and IE) of 1000 replicates with *r*_0_=0.1.

For simplicity, we assume that deleterious alleles anywhere on the genome have the same selective effects and dominance coefficients; this assumption is relaxed in Appendix D, SI. We use *Ks*_*c*_ to denote the critical selection strength per homozygous deleterious allele (scaled by the carrying capacity *K*), such that a non-zero equilibrium *N*_*_ exists for *Ks>Ks*_*c*_ but not for *Ks<Ks*_*c*_. Figure 1(a) shows *Ks*_*c*_ as a function of 2*LU* =2*L*(*u/r*_0_), the total mutation rate relative to the baseline growth rate, for different *Ku* (different colors) for nearly recessive (*h*=0.02; solid lines) and additive (*h*=0.5; dashed lines) alleles.

For populations to be metastable, the total genetic load must be less than the baseline growth rate. The total load scales with the mutation target *L*; the load per locus is ~2*u* for strongly deleterious variants (*u/hs*≪1 and *Ks*≫1). For smaller *Ks*, drift will typically inflate load above this deterministic expectation when deleterious alleles are additive (*h*~1/2) but may also reduce load (via more efficient purging) when alleles are recessive (*h*~0). This is reflected in figure 1(a): the threshold *Ks*_*c*_ required for metastable populations is higher for additive deleterious alleles (dashed lines) than recessive alleles (solid lines). Moreover, since genetic load in this drift-dominated regime depends only weakly on the mutational input *Ku*, the threshold *Ks*_*c*_ is largely independent of *Ku* (different colors). Finally, for all parameter combinations, the threshold *Ks*_*c*_ increases as the mutation target becomes larger: this simply reflects the fact that for the total load to be less than the baseline growth rate, the load per locus must be lower (requiring stronger selection) if deleterious variants segregate at a greater number of loci. Accordingly, for very large mutation targets 2*LU* =2*L*(*u/r*_0_)≳1, the total load will exceed *r*_0_ (and populations will fail), irrespective of the strength of selection against deleterious mutations.

The critical selection thresholds for metastability shown in fig. 1(a) are computed by neglecting demographic stochasticity, i.e., by assuming *ζ*=*r*_0_*K* to be very large. However, for moderate *ζ*, stochastic fluctuations in reproductive output from generation to generation may accelerate extinction: this effect can be especially significant in smaller populations as these tend to fix more deleterious alleles, which further reduces fitness and size, thus rendering populations even more vulnerable to stochastic extinction.

To investigate how demographic stochasticity contributes to extinction, we compare populations residing on islands with different carrying capacities *K*, but characterised by the same values of *Ks*, *Ku*, 2*LU*=2*L*(*u/r*_0_) and *h*. Mathematically, this involves taking the limit *s*→0, *u*→0, *K*→∞, *L*→∞, while holding *Ks*, *Ku*, 2*LU* constant: then, increasing *K* has the sole effect of weakening demographic stochasticity (by increasing *r*_0_*K*). Populations are initially perfectly fit, but accumulate deleterious variants over time, eventually becoming extinct due to the combined effects of genetic load and demographic stochasticity. Figure 1(b) shows the extinction ‘half-time’ *T*_1/2_=*r*_0_*t*_1/2_ (scaled by the baseline growth rate *r*_0_)— the time by which precisely half of all 1000 simulation replicates are extinct, as a function of 2*LU* for various *K* for nearly recessive (main plot) and additive alleles (inset). All results are from allele frequency simulations (assuming LE and zero inbreeding). Dashed vertical lines indicate the threshold 2*LU* above which metastable populations (with *N*_*_=1−*E*[*R*_*g*_|*N*_*_]>0) cannot exist (even for large *r*_0_*K*).

We find that extinction times increase with increasing *K* for all parameters. However, for parameter combinations that correspond to extinction in the large *r*_0_*K* limit (i.e., to the right of the dashed lines), this increase is approximately linear in *K*, while for parameters leading to metastability in the large *r*_0_*K* limit (left of dashed lines), this increase is faster than linear. In fact, in the metastable regime, even with a carrying capacity *K* of a few thousand individuals, demographic stochasticity will be sufficiently weak and extinction times large enough that isolated populations can persist over geological timescales: for these parameter regimes, it is environmental fluctuations (which our model ignores), rather than mutation load or demographic stochasticity, that will primarily influence extinction risk.

We also compute the average (scaled) population size *N*=*n/K* in the metastable state (fig. 1(c)). This declines with increasing 2*LU*, i.e., as we approach the threshold for loss of metastability, and is close to the semi-deterministic prediction (dashed black lines) for all values of *K*.

### Effect of migration on equilibrium population sizes and genetic load

We now consider how migration from a large mainland influences population dynamics on the island, for different genetic architectures of load, i.e., given certain selective effects and dominance coefficients of deleterious alleles. As before, we assume that all deleterious alleles have equal selective effects and dominance values, relegating the discussion of more general scenarios, where alleles with different fitness effects segregate, to the SI (Appendix D). We first identify critical parameter thresholds associated with qualitative changes in population outcomes for the two extremes: nearly recessive (*h*=0.02 in fig. 2) and additive (*h*=0.5) alleles. We then consider how these thresholds depend on the dominance of deleterious alleles (fig. 3).

**Figure 2:**
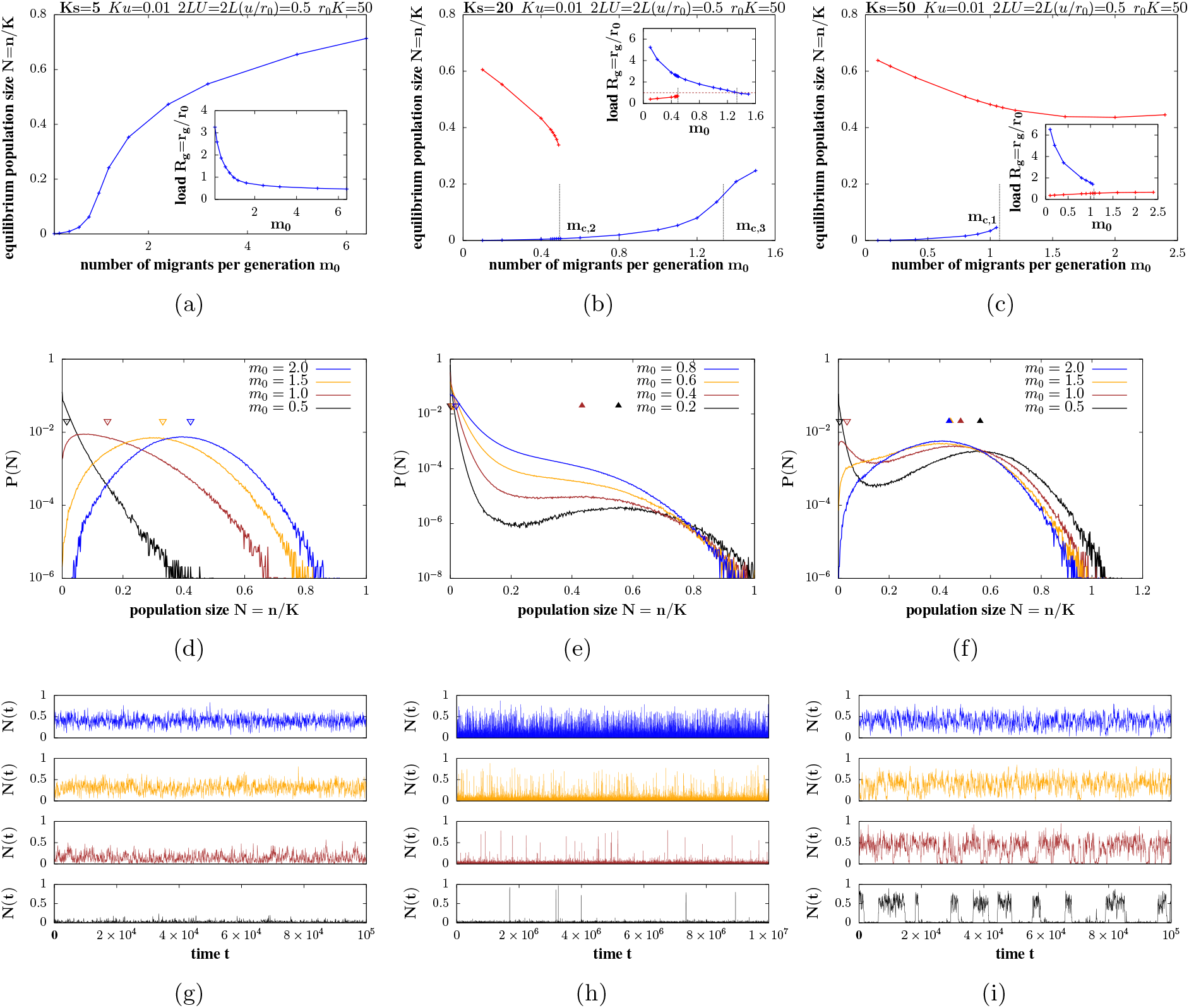
Effect of migration on equilibrium population sizes and load for weakly deleterious (*Ks<Ks*_*c*_; left column), moderately deleterious (*Ks*≳*Ks*_*c*_; middle) and strongly deleterious (*Ks≫Ks*_*c*_; right) nearly recessive (*h*=0.02) alleles. (A)-(C) Population size (main plots) and genetic load (inset), corresponding to one or more stable equilibria, vs. *m*_0_, the number of migrants per generation. Equilibria are obtained by numerically solving eq. (3) (semi-deterministic predictions). Blue lines represent the sink equilibrium (with an associated population size that tends to zero as *m*_0_→0); red lines represent the large-population equilibrium (with non-zero population size in the *m*_0_→0 limit). The thresholds *m*_*c,*2_ (2B) and *m*_*c,*1_ (2C) represent critical migration thresholds at which the sink or the large-population equilibrium vanishes. The threshold *m*_*c,*3_ (2B) is the critical migration level at which the (scaled) load associated with the sink state becomes less than 1. (D)-(F) Equilibrium probability distributions of the scaled population size *N*=*n/K* for various values of *m*_0_, as obtained from simulations (under LE and IE) in the three parameter regimes. The filled and empty triangles indicate the population sizes corresponding to alternative equilibria as predicted by the semi-deterministic analysis. (G)-(I) Time series *N*(*t*) vs. *t* over an arbitrary period after equilibration, for a single randomly chosen stochastic realisation in each of the three regimes. All plots show results for: *Ku*=0.01, 2*LU*=2*L*(*u/r*_0_)=0.5, *ζ*=*r*_0_*K*=50, *h*=0.02 and *Ks*=5, 20, 50 for the left, middle and right columns respectively (the critical threshold is *Ks*_*c*_~7.65). In addition, for the simulations (plots D-I), we use *r*_0_=0.1.

**Figure 3:**
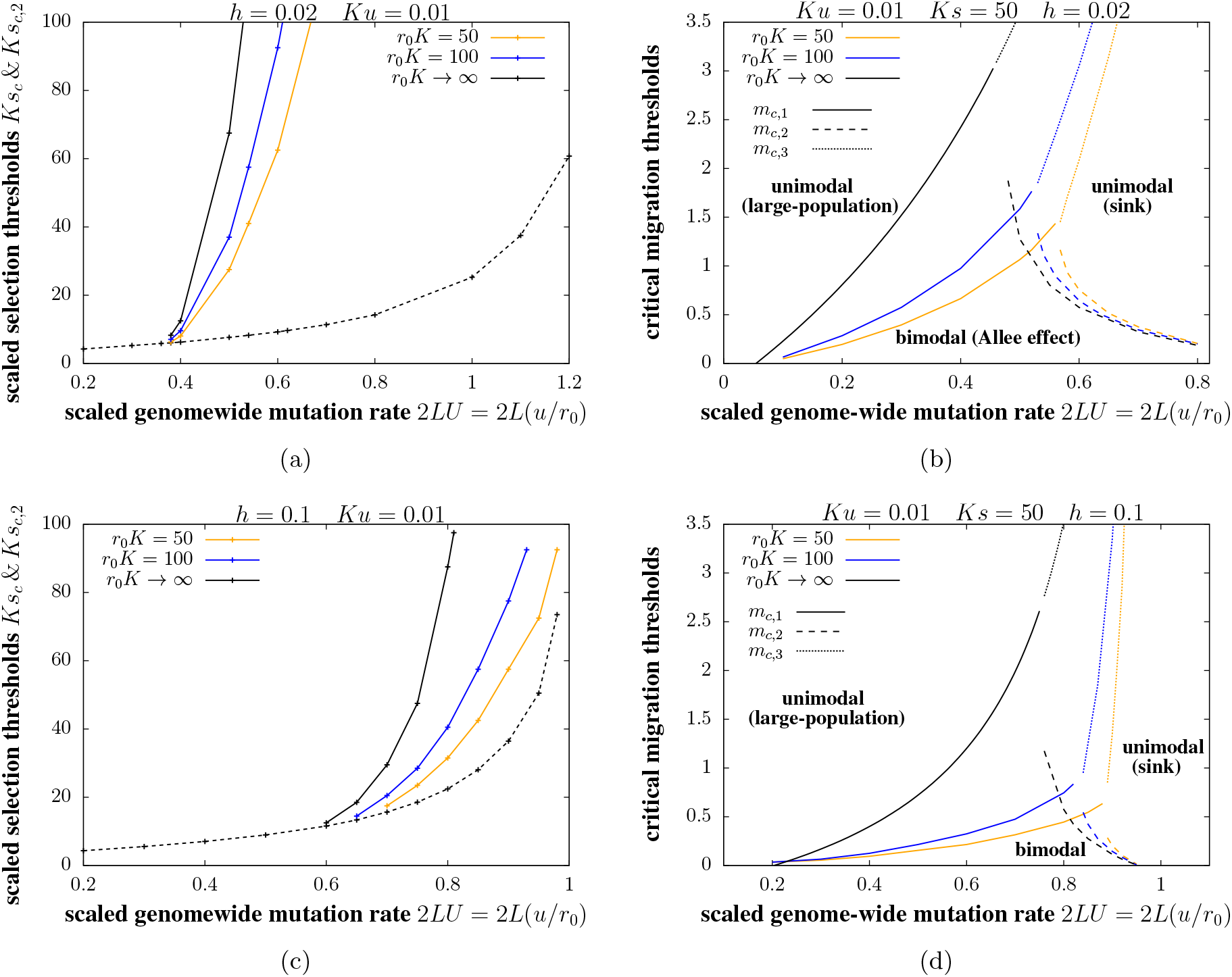
Semi-deterministic predictions for critical thresholds with and without accounting for demographic effects of migration. (A) and (C): Critical selection thresholds *Ks*_*c*_ (dashed line) and *Ks*_*c,*2_ (solid lines) as a function of the scaled mutation target 2*LU*=2*L*(*u/r*_0_) with (A) *h*=0.02 and (C) *h*=0.1, for *r*_0_*K*=50, *r*_0_*K*=100 and for *r*_0_*K*→∞ (in which limit migration has negligible demographic effects). The parameter *r*_0_*K*, which governs the magnitude of the demographic effect of migration (via the term *M*_0_=*m*_0_/(*r*_0_*K*)) is varied by changing *K*, while simultaneously varying *s*, *u* and *L* such that *Ks*, *Ku* and 2*LU*=2*L*(*u/r*_0_) are constant (so that the genetic effects of migration remain unchanged). The threshold *Ks*_*c*_ is such that populations rapidly go extinct for *Ks<Ks*_*c*_ in the limit of zero migration, but are metastable for *Ks>Ks*_*c*_, if starting from large but not small sizes (so that there is a genetic Allee effect). The second threshold *Ks*_*c,*2_ is such that increasing migration destabilizes the sink state for *Ks>Ks*_*c,*2_, but destabilizes the large-population state for *Ks*_*c*_<*Ks<Ks*_*c,*2_. The threshold *Ksc* is independent of migration, but *Ks*_*c,*2_ increases as *r*_0_*K* increases, i.e., as the demographic effect of migration becomes weaker. (B) and (D) The critical migration thresholds *m*_*c,*1_ (solid lines), *m*_*c,*2_ (dashed lines) and *m*_*c,*3_ (dotted lines) vs. 2*LU*=2*L*(*u/r*_0_) with (B) *h*=0.02 and (D) *h*=0.1, for various *r*_0_*K* for *Ks*=50. The threshold *m*_*c,*1_, which separates parameter regimes with bimodal population size distributions (characterised by alternation between the sink state and the large-population state) and unimodal size distributions (populations always in the large-population state) increases with increasing *r*_0_*K*. The threshold *m*_*c,*2_, which separates parameter regimes with bimodal size distributions and unimodal distributions (populations always in the sink state) decreases with increasing *r*_0_*K*. The threshold *m*_*c,*3_, which separates parameter regimes with load greater than or less than the baseline growth rate *r*_0_, increases with increasing *r*_0_*K*. All predictions are for *Ku*=0.01.

Figure 2 illustrates the effect of migration on population sizes and load on the island for three parameter combinations, corresponding to weakly deleterious (*Ks<Ks*_*c*_; left column), moderately deleterious (*Ks*≳*Ks*_*c*_; middle) and strongly deleterious (*Ks*≫*Ks*_*c*_; right) nearly recessive alleles (*h*=0.02). Recall that *Ks*_*c*_ is the selection threshold (obtained by neglecting demographic stochasticity) such that populations rapidly go extinct in the absence of migration for *Ks<Ks*_*c*_, but can be metastable (even without migration) for *Ks>Ks*_*c*_ and *r*_0_*K*→∞ (fig. 1).

Figures 2(a)-2(c) show population sizes corresponding to stable equilibria of eq. (3) vs. *m*_0_, the number of migrants per generation, for *Ks<Ks*_*c*_, *Ks*≳*Ks*_*c*_ and *Ks*≫*Ks*_*c*_. The insets show the equilibrium load vs. *m*_0_. Since these equilibria are calculated by numerically solving eq. (3), they represent *semi-deterministic* predictions, i.e., they neglect demographic stochasticity but account for the effects of drift, assuming that populations spend enough time in any metastable state that the distribution of allele frequencies can equilibrate. Figures 2(d)-2(f) show the stochastic distribution of the scaled population size *N*=*n/K*, as obtained from allele frequency simulations (assuming LE and zero inbreeding) after the population has equilibrated, for various *m*_0_, in the three parameter regimes. Figures 2(g)-2(i) show the corresponding time series *N*(*t*) over an arbitrary period, after equilibration, for a single randomly chosen stochastic realisation.

#### Weakly deleterious recessive alleles

For *Ks<Ks*_*c*_, there exists a single stable equilibrium at all migration levels (fig. 2(a)), with the corresponding population size approaching zero as migration becomes rarer. Accordingly, the stochastic distribution of *N* exhibits a single peak for all *m*_0_ (fig. 2(d)). An increase in migration causes the genetic load to decrease (inset, fig. 2(a)) and the equilibrium population size to increase. The stochastic distribution of population sizes is approximately exponentially distributed for low *m*_0_ (corresponding to a sink state), but shifts towards higher *N* as *m*_0_ increases, and is approximately normally distributed about the deterministic equilibrium *N*_*_ (indicated by empty triangles in fig. 2(d)) for large *m*_0_.

Increasing migration has two effects in this case: it reduces genetic load, typically by reducing homozygosity— a *genetic* effect, but also increases population numbers— a *demographic* effect. The genetic effect of migration, i.e., the dependence of the expected load 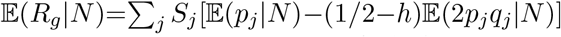 on *m*_0_, can be further decomposed into effects on the expected frequencies 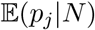 and expected heterozygosity 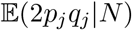 of deleterious alleles. For weakly deleterious recessive alleles, the expected frequency decreases with increasing migration for low *m*_0_, is minimum at intermediate *m*_0_, and then increases as *m*_0_ increases further (not shown). However, genetic load still decreases monotonically with increasing migration (inset, fig. 2(a)), due to the much sharper increase in heterozygosity with migration.

#### Strongly deleterious recessive alleles

For *Ks*≫*Ks*_*c*_, there exist *two* stable equilibria of the semi-deterministic population size dynamics (eq. (3)) at low levels of migration: population sizes and values of load corresponding to these two equilibria are shown in blue and red in fig. 2c). One equilibrium (blue lines in fig. 2(c)) corresponds to a sink state with the associated population size approaching zero as *m*_0_→0. The other equilibrium (depicted in red) corresponds to a large population, which can persist at finite fraction of carrying capacity even as migration becomes exceedingly rare, i.e., has size *N*_*_, which approaches 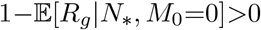 as *m*_0_→0.

The existence of two stable equilibria at low *m*_0_ translates into *bimodal* population size distributions (in black and brown in fig. 2(f)), with one peak close to extinction and the other at the semi-deterministic *N*_*_ (depicted by filled triangles). Population size *N*(*t*) also exhibits characteristic dynamics (see time series in black and brown in fig. 2(i)): populations tend to remain ‘stuck’ in either the sink state or the large-population state for long periods of time, typically exhibiting rather small fluctuations about the characteristic size associated with either state, and only transitioning rarely between states.

The sink and large-population states can also be thought of as “high-load” (i.e., genetic load *r*_*g*_ greater than the baseline growth rate *r*_0_) and “low-load” (*r*_*g*_<*r*_0_) states. Thus, in this parameter regime, populations exhibit a genetic Allee effect, wherein load is sufficiently low and net growth rates positive only above a threshold population size: therefore, starting from an initially empty island, populations cannot grow deterministically (and persist instead as migration-fed sinks), even though large populations can maintain themselves, at least in the absence of demographic fluctuations. Transitions between the sink (high-load) and large-population (low-load) states are thus inherently stochastic, arising due to demographic fluctuations (which are aided by higher migration) rather than due to a systematic drive towards the alternative state.

Changes in *m*_0_ have qualitatively different effects on the two states: at low *m*_0_, increasing migration reduces load and thus increases numbers in the sink state (blue curve), but has the *opposite* effect on the large-population equilibrium (red curves). This is due to the non-monotonic dependence of the frequency of deleterious recessive alleles on population size: increasing size causes drift to become weaker relative to selection but also reduces homozygosity, so that fewer deleterious alleles are exposed to selection; frequencies are thus minimum at intermediate population sizes, reflecting the tension between these two opposing effects. In small sink populations, migration from the continent reduces homozygosity as well as deleterious allele frequencies, thus reducing load. However, in larger populations, increasing migration reduces the homozygosity but raises the frequency of deleterious alleles (since deleterious alleles are purged less efficiently in very large mainland populations than in intermediate-sized or large island populations); the overall effect of migration is thus to increase load in the large-population state.

Above a critical migration threshold, which we denote by *m*_*c,*1_, the sink equilibrium vanishes. Thus, for *m>m*_*c,*1_, populations always have a positive growth rate and reach a finite fraction of carrying capacity, regardless of starting size. As before, this qualitative change in the (semi-deterministic) population size dynamics at a critical migration level has its analog in a qualitative change in the stochastic distribution of population sizes (fig. 2(f)): increasing migration causes the population size distribution to change from bimodal to unimodal (with a sole peak at the large-population equilibrium 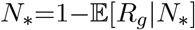). At higher migration rates, i.e., as we approach to the critical migration threshold *m*_*c,*1_ starting from low *m*_0_, turnover between the sink and large-population states becomes more rapid— note the shorter intervals between transitions in the brown vs. black time series in fig. 2(i).

#### Moderately deleterious recessive alleles

Consider now the case where selection is stronger than the threshold *Ks*_*c*_ but not too strong, i.e., *Ks*≳*Ks*_*c*_ (middle column in fig. 2). As in the *Ks*≫*Ks*_*c*_ case, there exist two alternative stable equilibria at low migration levels, one corresponding to the sink state (blue lines in fig. 2(b)) and the other to the large-population state (red lines). As before, increasing migration alleviates inbreeding load and increases population size in the sink state but elevates load (by hindering purging) and decreases population size in the large-population state. Unlike in the *Ks*≫*Ks*_*c*_ state, the latter effect is much stronger: thus, above a critical migration threshold, which we denote by *m*_*c,*2_, it is the large-population equilibrium that vanishes, so that there is only a single (sink) equilibrium for *m>m*_*c,*2_.

The analogous change in the population size distribution with increasing migration (fig. 2(e)) is somewhat subtle: for the lowest value of *m*_0_ (black), the distribution is clearly bimodal, with most of the weight close to *N*=0 (sink state) and a very small peak at the large-population equilibrium. As migration increases, the distribution of sizes associated with the sink state widens, while the peak corresponding to the large-population equilibrium shifts towards lower *N* (e.g., brown plot). At high migration levels (orange and blue plots), there is no distinct second peak and only the sink state persists. In this state, the genetic load exceeds *r*_0_ *on average*; however, its distribution and the corresponding distribution of *N* is quite wide.

If migration increases further, then the average load associated with the sink state continues to fall; at a third threshold (denoted by *m*_*c,*3_; here ~1.33), the load again becomes lower than the baseline growth rate *r*_0_ or, alternatively, scaled load *R*_*g*_=*r*_*g*_/*r*_0_ less than 1 (depicted by a dashed line in fig. 2(b)). Thus, for *m*_0_*>m*_*c,*3_, populations can grow, starting from small numbers, and reach a finite fraction of carrying capacity. However, unlike the large-population state that emerges at higher *Ks*, in this case, populations are highly dependent on migration and would rapidly collapse (due to the fixation of deleterious alleles) if cut off from the mainland.

Population size dynamics (fig. 2(h)) are characterised, as in the *Ks*≫*Ks*_*c*_ case, by occasional transitions between the sink state and the large-population state for low values of *m*_0_ (black), with transitions becoming more frequent with increasing *m*_0_ (orange). Transition times are much longer than in the *Ks*≫*Ks*_*c*_ case (note the values on the x-axis); thus transitions are unlikely to occur over realistic timescales, and populations will typically be observed in the sink state. At higher migration levels, there are no obvious transitions, with population sizes and load fluctuating (with some skew) about a mean value.

#### Additive alleles

With additive effects (*h*~0.5), any deleterious allele experiences the same selective disadvantage, irrespective of whether it appears in the heterozygous or homozygous state: thus, there is no purging in smaller populations (which have higher homozygosity) and allele frequencies decrease monotonically with increasing population size. As a result, migration from the larger mainland always decreases load (by decreasing the deleterious allele frequency) and increases the size of the island population.

This implies that there are only two qualitatively distinct regimes: with weakly deleterious alleles (*Ks* less than the corresponding *Ks*_*c*_; see fig. 1(a)), populations tend to extinction as migration declines. There is a single equilibrium of the semi-deterministic population size dynamics for all *m*_0_ or analogously, a single peak of the stochastic population size distribution, which shifts towards higher sizes as *m*_0_ increases.

For strongly deleterious alleles (*Ks>Ks*_*c*_), population size *N* and load *R*_*g*_ are largely insensitive to migration, since populations can always grow, starting from small numbers, at least when demographic stochasticity is unimportant (i.e., for *r*_0_*K*≫1). With smaller *r*_0_*K*, population sizes and load depend weakly on migration, since demographic stochasticity may depress *N* and inflate the effects of drift (even with *Ks>Ks*_*c*_). Moreover, in this regime, populations are also prone to stochastic extinction for *m*_0_<1/2 [14], such that the distribution of *N* is inherently bimodal. Thus, where demographic stochasticity is significant, a low level of migration may be needed for stable populations even with strong selection against additive alleles.

The analysis outlined here (based on eq. (3)) also applies to loci with a distribution of effects: in Appendix D (SI), we consider examples where a fraction of deleterious mutations are additive and the remaining fraction recessive. We also compare the results of allele frequency simulations with those of individual-based simulations with unlinked loci (Appendix C, SI). These show fairly close agreement, suggesting that LD and ID do not significantly affect allele frequency dynamics, at least for the typical parameter values considered here: this is also consistent with earlier work, which suggests only a modest effect of disequilibria on background selection in sub-divided populations under soft selection [19].

### Disentangling genetic vs. demographic effects of migration on recessive alleles

In summary, given a mutation target 2*LU*≲1 and assuming equal-effect loci, there is a critical selection threshold *Ks*_*c*_, such that large populations are metastable in the *m*_0_→0 limit for *Ks> Ks*_*c*_ but not for *Ks<Ks*_*c*_. For *Ks<Ks*_*c*_, there is a genetic Allee effect at low migration levels and with partially recessive alleles: populations cannot grow subsequent to recolonisation of an initially empty island, and persist only as demographic sinks until a chance fluctuation increases numbers sufficiently that load can be purged and the alternative (large-population) equilibrium attained; such large populations can then be maintained over long periods of time (fig. 2(i)). For very low dominance values (*h*≲0.15), there is a second threshold *Ks*_*c,*2_ with *Ks*_*c,*2_*>Ks*_*c*_ (see below), which separates parameter regimes characterised by qualitatively different effects of migration on population outcomes: for *Ks>Ks*_*c,*2_, increasing migration destabilizes the sink state, so that only the large-population state persists above a migration threshold *m*_*c,*1_, whereas for *Ks*_*c*_<*Ks<Ks*_*c,*2_, increasing migration destabilizes the large-population state, so that only the sink state persists above a threshold *m*_*c,*2_. However, such migration-fed sink populations can be quite large: above a third threshold *m*_*c,*3_, populations may even have sufficient heterozygosity to again attain a low-load (*r*_*g*_<*r*_0_) state.

To what extent can we attribute such qualitative changes in population outcomes to the genetic vs. demographic effects of migration? As before, one approach is to compare critical thresholds for populations with the same scaled parameters *Ks*, *Ku*, 2*LU*=2*L*(*u/r*_0_) and *h*, while increasing the carrying capacity *K* (simultaneously increasing *L* and lowering *u*, *s*). Then, as *K* increases, the demographic effects of migration (which depend on the dimensionless parameter *M*_0_=*m*_0_/(*r*_0_*K*); see eq. (1)) can be neglected, while its genetic effects on load (which depend on *Ks*, *Ku* and *m*_0_) remain important.

Figure 3 shows these comparisons for two dominance values— *h*=0.02 (figs. 3(a), 3(b)) and *h*=0.1 (figs. 3(c), 3(d)). Note that we only consider predictions of the semi-deterministic analysis (eq. (3)), which assumes that population sizes are sharply clustered around the peaks of the distribution and transitions between peaks are infrequent. This assumption clearly breaks down close to transition thresholds (e.g., see figs. 2(e), 2(f)); thus, thresholds observed in simulations may differ somewhat from semi-deterministic predictions (details in Appendix B, SI). Moreover, the semi-deterministic analysis neglects all demographic stochasticity, and thus does not account for the fact that changes in *K* will affect not just the (systematic) demographic effect of migration but also the (stochastic) effect of demographic fluctuations. Nevertheless, the semi-deterministic analysis is useful as it allows us to explore qualitative dependencies of critical thresholds on the underlying parameters without resorting to time-consuming simulations.

Figures 3(a) and 3(c) show the critical selection thresholds *Ks*_*c*_ (as in fig. 1(a)) and *Ks*_*c,*2_ vs. the scaled mutation target 2*LU*=2*L*(*u/r*_0_) for the two values of *h*, for various carrying capacities (with *Ks*, *Ku*, 2*LU*=2*L*(*u/r*_0_) held constant, as *K* is varied). The threshold *Ks*_*c*_, which relates to population outcomes in the zero migration limit, is (by definition) independent of *K* in the semi-deterministic setting, as increasing *K* merely weakens the demographic effects of migration. The threshold *Ks*_*c,*2_ decreases as *K* decreases, i.e., as the demographic effects of migration become stronger. This means that when alleles are moderately deleterious, i.e., characterised by a certain intermediate value of *Ks*, populations are stabilized (i.e., rescued from recurrent collapse into the high-load sink state) by increasing migration more easily on *smaller* islands, where the demographic effects of migration are stronger.

Figure 3(b) and 3(d) show the critical migration thresholds *m*_*c,*1_, *m*_*c,*2_ and *m*_*c,*3_ vs. 2*LU* for a given *Ks* value (chosen to be 50). Here, with *h*=0.02 for example (fig. 3(b)), increasing migration causes the sink state to vanish for 2*LU*≲0.5, but degrades the large-population (low-load) state for 2*LU*≳0.5. The threshold *m*_*c,*1_ (solid lines) is highly sensitive to *K*: populations can be stabilized at a finite fraction of carrying capacity at much lower levels of migration on smaller islands (given *Ks*, *Ku*), suggesting a key role of the demographic effects of migration. We can obtain an explicit expression for the threshold *m*_*c,*1_ in the limit *K*→∞ (so that *M*_0_= *m*_0_/(*r*_0_*K*), which governs the demographic effects of migration, is negligible):

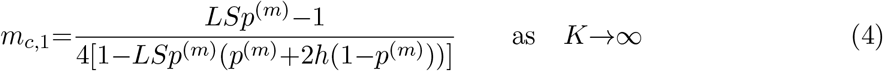

Interestingly, this threshold depends only on the load, *LSp*^(*m*)^(*p*^(*m*)^+2*h*(1−*p*^(*m*)^)), and breeding value, *LSp*^(*m*)^, among migrants, and is independent of *Ks* and *Ku*: this simply reflects the fact that the growth rate of a very small population (just after recolonisation) depends primarily on the genetic composition of founders (and not selection on their subsequent descendants). Thus, the critical level of migration required to prevent a genetic Allee effect in the limit of very large carrying capacities is also independent of *Ks* and *Ku*.

The threshold *m*_*c,*2_, which signals the collapse of the large-population state, is less sensitive to *K*: this is consistent with our expectation that demographic effects of migration should be less important when numbers are larger. There is, nevertheless, a moderate decrease in *m*_*c,*2_ with *K*, which can be rationalised as follows: the (detrimental) genetic effects of migration on the large-population state are more effectively compensated by its demographic effects when islands are smaller (lower carrying capacity), allowing the large-population state to persist despite higher levels of migration on such islands. Finally, the third threshold *m*_*c,*3_, which signals the emergence of a migration-dependent low-load state (at large 2*LU*) is also highly sensitive to the demographic effects of migration, and becomes unrealistically high in the large *K* limit, where increasing migration can only shift allele frequencies but has little effect on population numbers (relative to carrying capacity).

A comparison of the top and bottom rows in fig. 3, which correspond respectively to *h*=0.02 and *h*=0.1, shows that as deleterious alleles become less recessive (larger *h*), the parameter regime in which increasing migration eliminates the large-population equilibrium shrinks drastically, and only emerges for very high genome-wide mutation rates — with *h*=0.1, the second threshold *Ks*_*c,*2_ exists only for 2*LU*≳0.8 in fig. 3(c). This is consistent with the fact that purging of recessive mutations in smaller populations (which allows them to evolve significantly lower load than mainland populations) is only effective for very low *h*; thus, gene flow from the mainland is less detrimental to the large-population equilibrium in island populations for less recessive alleles. In fact, for *h*≳0.15, increasing migration always causes the sink equilibrium to vanish, irrespective of *Ks*. Moreover, the migration threshold *m*_*c,*1_ at which the sink state vanishes falls with increasing *h* (compare figs. 3(b) and 3(d)). Thus, we observe a genetic Allee effect only at very low migration rates for moderately recessive alleles. This is consistent with the fact that inbreeding load in small populations (due to excess homozygosity) becomes lower as alleles becomes less recessive, and is alleviated by even low levels of migration.

## Discussion

A key parameter governing the fate of peripheral populations is 2*LU*=2*L*(*u/r*_0_), the genome-wide deleterious mutation rate relative to the baseline rate of population growth: low-load (large-population) states are possible only for 2*LU*≲1, provided selection against deleterious variants is sufficiently strong and/or migration high. Conversely, for 2*LU*≳1, populations exist only as demographic sinks, irrespective of selection strength. The parameter 2*LU* is a measure of the ‘hardness’ of selection and can be small either if the total mutation rate 2*Lu* (which determines total load in the absence of drift) is small or, more realistically, if the growth rate *r*_0_ i.e., the logarithm of the baseline fecundity, is high (corresponding to the soft selection limit).

Our analysis highlights qualitatively different effects of migration on population outcomes, depending on fitness effects and the total mutation target of deleterious variants (figs. 2 and 3). For example, with 2*LU*=0.5 (which corresponds to a ≈50% reduction in growth rate due to genetic load in a deterministic population), typical *Ks* must be at least ~5 for recessive and ~10 for additive alleles, if populations are to be metastable in the absence of migration. For weaker selection, gene flow from the mainland is beneficial, and aids population survival by hindering the fixation of deleterious alleles. However, for stronger selection and with recessive alleles, the fitness and size of ‘low-load’ island populations actually declines with increasing migration from the mainland (due to higher deleterious allele frequencies in the latter). In the most extreme scenario, where load is primarily due to segregation of recessive alleles of moderate effect, intermediate levels of migration may actually increase load so much that populations degenerate into high-load demographic sinks (fig. 2(b) and fig. S4 in SI).

We identify two regimes in which peripheral populations can maintain stable numbers at a substantial fraction of carrying capacity, with qualitatively different roles of migration in the two. When selection is strong, i.e., *Ks*≫*Ks*_*c*_ (for additive alleles) or *Ks>Ks*_*c,*2_ (for recessive alleles) for a given 2*LU*, genetic load is low and populations stable, largely independently of migration. In this case, low levels of migration (typically ≲1 migrant per generation; note typical values of *m*_*c,*1_ in fig. 3) are sufficient to prevent a genetic Allee effect, should demographic stochasticity or chance fluctuations in load drive population numbers down. On the other hand, with weakly or moderately deleterious alleles, stable populations rely on rather high levels of migration (≫1 migrant per generation; note typical *m*_*c,*3_ in fig. 3) and are only weakly differentiated with respect to the mainland. Here, migration is essential for maintaining heterozygosity and preventing fixation of deleterious alleles, even though numbers are relatively large.

Both classical quantitative genetics and analyses of allele frequency spectra suggest that most mutation load is due to weakly deleterious alleles [24, 25]. The weak selection on deleterious mutations may be strong relative to random drift in the species as a whole [24], but is likely to be dominated by random drift within local demes (i.e., *Ks<*1 in our notation). If this is so, then extinction can be avoided only if many migrants enter demes in each generation. Fortunately, this is generally the case: *F*_*ST*_ is typically small, implying that *m*_0_ is between 0.5 and 5 ([26]; note that *m*_0_ is the number of migrant genes, corresponding to 2Nm in the usual diploid notation). Thus, whilst selection can act effectively to suppress the mutation load in a well-connected metapopulation, demes that receive few migrants will be vulnerable to the accumulation of load due to weakly selected mutations. Moreover, when fitness has additional environment-dependent components, local adaptation must depend on alleles of intermediate effect and is hindered by high migration, especially for marginal habitats within the metapopulation [14].

What implications might our work have for the conservation of natural populations? Provided that several migrants are exchanged per generation, selection against deleterious alleles can be effective across the whole population. Indeed, subdivision into small subpopulations can help purge deleterious recessives, making selection more effective than with panmixia. Thus, random drift would lead to a severe load only if local demes are highly isolated - in which case environmental fluctuations are more likely to cause extinction than the gradual accumulation of weakly deleterious mutations [9]. Our work implies that an intermediate rate of migration minimises mutation load, by preventing extinction of local populations, and yet still allowing some purging. However, the extinction risk arising from environmental fluctuations (which we underestimate by including only demographic stochasticity) favours higher migration. Conversely, local adaptations require selection that is stronger than drift within local demes (*Ks>*1) [14]; if this is a concern, then substantial deme sizes are required in the long term.

Our model make various assumptions: first, we take mutation rates to and from the deleterious state to be equal. Asymmetry in mutation rates would not qualitatively alter our conclusions as long as there is even weak migration (*m*_0_≳0.1), as load is then alleviated primarily by migration rather than reverse mutation. However, with zero migration and *no* reverse mutation, selection must be strong enough to prevent long-term ratchet-like accumulation of deleterious variants, resulting in a much higher threshold *Ks*_*c*_ for metastability.

Second, we assume deleterious allele frequencies on the mainland to be close to deterministic. This assumption is not crucial: our qualitative conclusions remain unaltered as long as mainland frequencies are much lower than typical island frequencies. However, if mainland populations are small enough to harbour deleterious alleles at high frequencies at a subset of loci (which would, in general, be different from the loci fixed for deleterious alleles on the island), then we expect heterosis and the beneficial effects of migration to be weaker (with one-way migration between the mainland and island). More generally, extending this analysis to the co-evolution of load and population sizes in a metapopulation, where each sub-population may be close to fixation for different deleterious alleles, remains an interesting direction for future work.

Third, we assume a rather simple genetic architecture of load: loci are assumed to be unlinked, and load additive across loci. Deviations from additivity, e.g., synergistic epistasis between deleterious variants can lower load [30, 31], and may arise, for example, if multiple traits are under stabilizing selection. However, linkage between deleterious variants can inflate load by compromising selection efficacy at individual loci via Hill-Robertson interference [27], making it difficult to arrive at general predictions for the effects of selective interference (due to linkage and epistasis between selected variants) on the eco-evolutionary dynamics of marginal populations. Fourth, we ignore environmental stochasticity and demographic Allee effects, which may strongly influence outcomes [8, 32], especially in parameter regimes where mutation accumulation and demographic stochasticity in themselves are unlikely to cause extinction.

Finally, our analysis relies on a semi-deterministic analysis, which accounts for genetic drift but neglects demographic stochasticity. While this approximation captures qualitatively different population states across parameter space, it gives little insight into the dynamics. In particular, where alternative, i.e., low-load and high-load states are possible, the key assumption underlying the semi-deterministic analysis — namely, that allele frequencies have sufficient time to equilibrate at any given population size— is satisfied only when populations are in one or other state, and not while they transition between states. This makes it challenging to describe the complex co-evolution of load and population size during transitions and arrive at a complete understanding of transition timescales.

Our aim has been to base our analysis on as few parameters as possible, in the hope that these can be related to observations from nature. We have reasonably good estimates of the fitness effects of deleterious mutations, and their degree of dominance - albeit largely from Drosophila [24]. We also now have accurate measures of the total mutation rate; the total rate of deleterious mutation is still uncertain [28], but may be substantial in complex organisms [29]. Population structure is less well understood: we have very many estimates of *F*_*ST*_ [26], which reflects the numbers of incoming migrants, but local effective deme size is harder to estimate, even if demes can be defined at all. However, the common observation of heterosis implies that different deleterious recessives are common in different populations, suggesting a substantial drift load.

The rather complex effects of migration observed even in this relatively simple model with unconditionally deleterious alleles suggest that a comprehensive understanding of the effects of gene flow on eco-evolutionary dynamics at range limits must account for both environment-dependent (local) and environment-independent (global) components of fitness. These may be influenced by (partially) overlapping sets of genetic variants, so that genetic load is shaped fundamentally by pleiotropic constraints. Such extended models are key to understanding when, for example, assisted gene flow is beneficial, and whether its mitigatory effect on inbreeding depression may be outweighed by any outbreeding depression that it might generate.

From a conceptual viewpoint, our analysis highlights the importance of considering explicit population dynamics when analysing the influence of gene flow on the efficacy of selection in sub-divided populations. Simple predictions, e.g., that a sub-divided population under hard selection should behave as a single population with an inbreeding coefficient equal to *F*_*ST*_ [33, 18], may break down when sub-populations can undergo local extinction. In this case, purging may be ineffective and the efficacy of selection reduced relative to undivided populations, in contrast to standard predictions that subdivision always reduces load when selection is hard. We regard the framework proposed here as a starting point for detailed studies in specific metapopulations, which take into account the joint evolution of population size and mutation load.

## SUPPLEMENTARY INFORMATION

### A. Description of simulations

We carry out two kinds of simulations: simulations assuming LE (linkage equilibrium) and zero inbreeding, which only track allele frequencies at the *L* loci and population size *N* as a function of time; and individual-based simulations, which make no simplifying assumptions and track whole diploid genomes (with *L* loci) of all individuals present in the island population.

#### Simulations assuming LE and zero inbreeding

If recombination is faster than all ecological and evolutionary processes, then statistical associations between allelic states at different loci (linkage disequilibria or LD) can be neglected. In addition, if there is no significant inbreeding, then the probability of identity by descent at a locus, and correlations between identity by descent across loci (identity disequilibria or ID) are also negligible. Then, individual genotypes are simply random assortments of deleterious and wildtype alleles, and can be generated by independently assigning alternative allelic states to different loci with probabilities equal to the allele frequencies, allowing us to only track allele frequencies (instead of genotypes).

We initiate simulations by assuming that there are *K* individuals on the island and that the deleterious allele at each locus is absent. We then evolve the population in discrete time by updating allele frequencies and population size in each generation to reflect the effects of migration, mutation, selection and reproduction, and genetic drift. Starting with frequencies {*p*_*j*_(*t*)} and size *n*(*t*) at the end of generation *t*, we first implement migration in generation *t*+1 by sampling the number of migrants *m* from a Poisson distribution with mean *m*_0_. The population size and allele frequencies are then updated as: 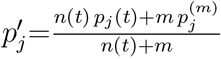 and *n*′=*n*(*t*)+*m*. Here, 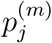 are the corresponding frequencies on the mainland.

Mutation has no effect on population size: *n*′′=*n*′; allele frequencies are changed as: 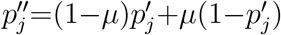.

The effects of selection and reproduction are then captured by changing allele frequencies according to: 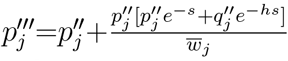, where 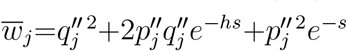 and 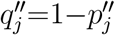. This is the standard equation relating allele frequencies before and after selection; it assumes that loci evolve independently (no indirect selection due to LD and ID), and that there is no inbreeding. The new population size *n*′′′ after selection and reproduction is generated by drawing a Poisson-distributed random variable with mean 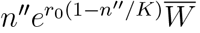, where the mean population fitness 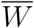 is calculated by multiplying the marginal mean fitnesses of all loci: 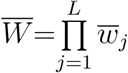. Note that sampling the population size from a Poisson distribution (rather than simply choosing it to be equal to the mean of the distribution) introduces demographic stochasticity into the simulation.

The new population size at the end of generation *t*+1 is just: *n*(*t*+1)=*n*′′′. The new allele frequencies {*p*_*j*_(*t*+1)} are generated by drawing Binomially distributed random variables {*X*_*j*_} with corresponding parameters *n*(*t*+1) and 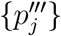 and then setting {*p*_*j*_(*t*+ 1) = *X*_*j*_/*n*(*t*+1). This last step, which amounts to standard Wright-Fisher sampling of the new allele frequencies based on the new population size *n*(*t*+1) and the deterministic allele frequencies 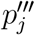, ‘adds’ the random effects of genetic drift on to the systematic effects of migration, mutation and selection (which are already captured by 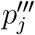).

The procedure described above updates {*p*_*j*_} and *n* over discrete generations by implementing the effects of migration, mutation, selection and drift in each generation sequentially. An alternative would be to numerically integrate the continuous time equations (eq. (1) in the main text). This is, however, prone to numerical error, and is computationally far more intensive. Both procedures are expected to yield very similar results if *m*_0_*/n, μ, s, r*_0_≪1.

While the allele frequency simulations described here are fast, a fundamental limitation is that the underlying assumptions (i.e., no significant inbreeding, and effects of drift and migration weaker than those of recombination) must break down close to extinction thresholds. Thus, we also carry out individual-based simulations (described below), that make no such assumptions and capture the evolution of all multi-locus associations.

#### Individual-based simulations

In this bottom-up simulation approach, we track eco-evolutionary dynamics in peripheral populations by explicitly following individuals, recording their allelic states and population numbers from one generation to another. Besides the advantage that comes from the inclusion of individual variations and interactions, this modeling framework allows us track the evolution of each locus as well as all the associations among loci (i.e. LD and ID).

We simulate a sexually reproducing population with random mating and non-overlapping discrete generations. At the start of the simulation, the island population is assumed to consist of *N* diploid individuals where the fitness of each individual is determined by *l* loci with two possible allelic state per locus coded by 1’s and 0’s. The 1 allele is considered the wildtype and the 0 allele the recessive deleterious variant so that the fitness of the three possible genotypes (11, 10 and 00) at each locus are respectively in the ratio 1:1−*hs*:1−*s* where *s* is the homozygous selective effect and *h* is the dominance coefficient affecting the expression of the deleterious allele. We assume multiplicative fitness across loci (i.e. no epistasis) so that the fitness of an individual with *n* heterozygous loci, *m* homozygous loci for the deleterious allele and *m*′ homozygous loci for the wild type allele can be expressed as (1−*hs*)^*n*^(1−*s*)^*m*^ where *n*+*m*+*m*′=*l*. It is worth noting that this fitness function is slightly different from that used in the main text, but the two approach each other in the limit of small *s* (assumed throughout the paper). The simulation is initialized by assuming that individuals are initially perfectly fit (i.e are composed of only the 0 allelic state) so that the frequency of the deleterious variant at each locus is 0. The order of events in the life cycle of individuals in each generation is taken as, mutation → migration → reproduction (meiosis and genetic recombination)+ density-dependent survival. These would be explained in a little more detail below.

Mutation: We model both forward and backward mutation rates i.e. with probability *μ*, there is a shift from an allelic state 1 to 0 and with probability *ν* there is a shift in the opposite direction. In this work, we assumed *μ*=*ν* although this is not true in general.

Migration: the migration step is implemented by choosing a random number, of individuals (drawn from a Poisson distribution with rate *m*_0_) to be migrants from a fixed mainland pool assumed to be in deterministic mutation-selection balance. Allelic states are then randomly assigned to each migrant locus such that the frequency of the deleterious variants (among migrants) at each locus are close to the deterministic predictions for a single locus under mutation-selection balance. Following migration, both the frequency at each locus as well as the population size on the island change.

Reproduction and density dependent survival: This phase of the simulation involves gamete formation (through the process of meiosis and genetic recombination) as well as a decision as to how many offspring survive to be parents in the next generation. The latter is done by sampling an integer number *l* from a Poisson distribution with rate 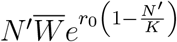 where *K* is the carrying capacity of the population, *N*′ is the population size after migration, 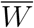 is the mean fitness of the population and *r*_0_ the population growth rate. Once this is determined, *l* pairs of parents are then selected (with replacement) from the population in proportion to their fitness to form offspring individuals. Each offspring gamete is formed by assuming that the allelic state contributed to the gamete at any given locus is sampled from the same or alternative chromosome in both parents with probability 0.5 (i.e. free recombination).

### B. Semi-deterministic approximation

The semi-deterministic approximation outlined in the main text applies when the distribution of population sizes is sharply peaked around one or more values. For a given set of parameters *Ks*, *Ku*, *h*, 2*LU*=2*L*(*u/r*_0_), and *m*_0_, we expect the peaks of the distribution to become sharper and the predictions of the semi-deterministic approximation to become more accurate for larger *r*_0_*K* (corresponding to weaker demographic fluctuations) and larger *L* (corresponding to weaker fluctuations in genetic load). Figure S1a illustrates this by plotting the distribution *P*(*N*) (as obtained from allele frequency simulations) for various values of *r*_0_*K* (which is changed by changing *K*). Note that in increasing *K*, while holding *Ks*, *Ku* and 2*L*(*u/r*_0_) constant, we must simultaneously lower *s* and *u*, while increasing *L* proportionately.

**Figure S1:**
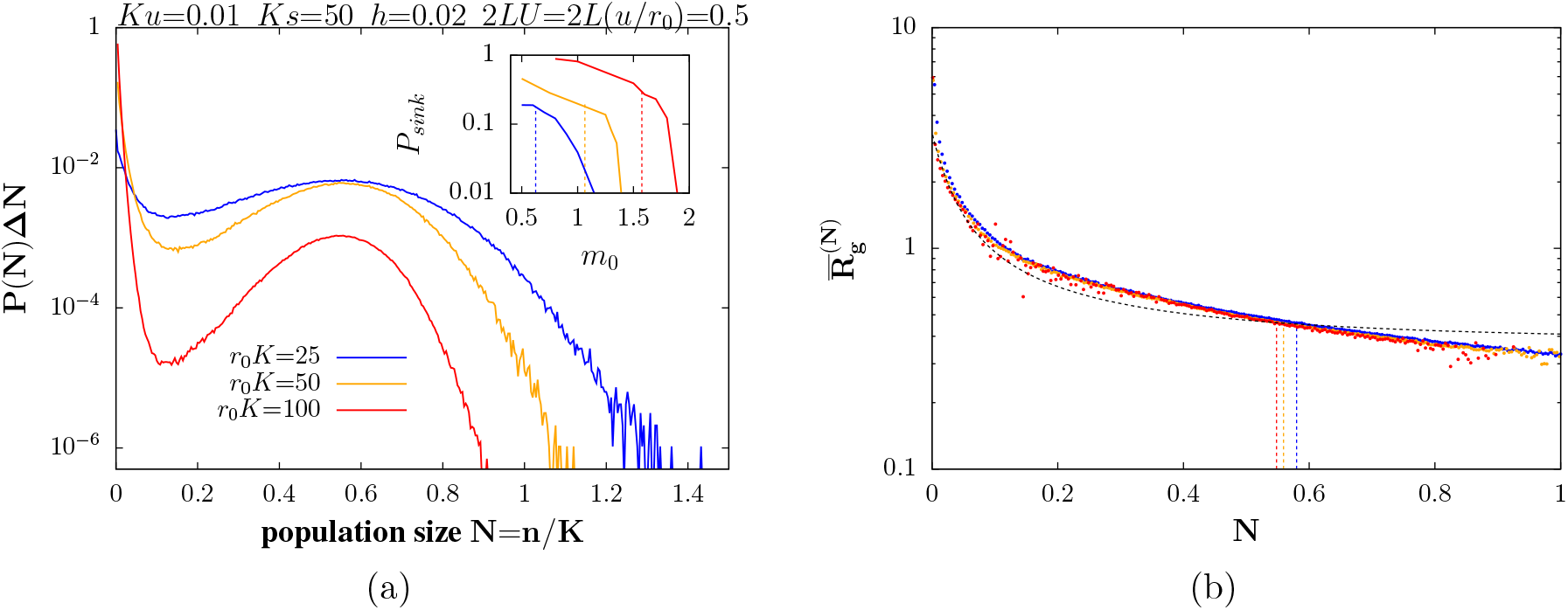
(A) Distribution of population sizes *P*(*N*) integrated over intervals of size Δ*N*=0.004 for *r*_0_*K*=25, 50, 100 (blue, orange, red), where *r*_0_*K* is varied by changing *K* (along with *s*, *u*, *L*), while keeping *Ku*, *Ks*, 2*LU*=2*L*(*u/r*_0_) and *r*_0_ constant. Inset: *P*_*sink*_, the probability that the population is in the sink state, vs. *m*_0_, the number of migrants per generation. Vertical dashed lines represent semi-deterministic predictions for the critical threshold *m*_*c,*1_, at which the sink equilibrium vanishes. *P*_*sink*_ is calculated by integrating the distribution *P*(*N*) from 0 to the minimum of the distribution (which lies between the sink and large-population equilibrium). (B) Average load 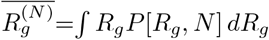 at a given *N* (points; different colours correspond to different *r*_0_*K*) and the expected load under mutation-selection-drift equilibrium for that *N*(dashed line) as a function of *N*. Average load equals the equilibrium expectation at the predicted large-population equilibrium (indicated by vertical dashed lines) for all *r*_0_*K*; the two quantities also match closely near *N*=0 for larger values of *r*_0_*K*. All results (solid lines in A and points in B) are from allele frequency simulations; dashed lines indicate various semi-deterministic predictions (as described above). Parameter values: *Ku*=0.01, *Ks*=50, *h*=0.02, 2*LU*=2*L*(*u/r*_0_)=0.5 and *r*_0_=0.1.

As expected, the peak corresponding to the large-population equilibrium becomes narrower, i.e., typical fluctuations in *N* about the average (in that state) become smaller, as *r*_0_*K* increases. The distribution of population sizes in the sink state also becomes more sharply clustered around 0, and the valley separating the large-population equilibrium from the sink state becomes deeper as *r*_0_*K* increases.

We then ask whether the basic approximation underlying the semi-deterministic analysis– that genetic load is close to its equilibrium expectation under drift-mutation-migration-selection balance when populations are in one or other metastable state– becomes increasingly accurate for larger *r*_0_*K*. Figure S1b shows the average load 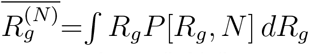 at a given *N* (as obtained from allele frequency simulations), as a function of *N* for various *r*_0_*K*. The average load (points) is different from the load expected under mutation-selection-drift-migration balance 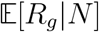 (black line) except under two conditions: first, the average load equals the equilibrium expectation near the large-population equilibrium (indicated by vertical dashed lines) for all values of *r*_0_*K*(different colours); second, the average load also matches the equilibrium expectation in the vicinity of *N*=0 for large *r*_0_*K*.

These observations are consistent with the general expectation that allele frequencies and genetic load will equilibrate only if population sizes remain roughly constant over the time scales required to reach mutation-selection-drift-migration balance: this condition can only be satisfied close to the peaks of the distribution (which correspond to equilibria of the semi-deterministic population size dynamics) and will typically not hold while populations transition between equilibria. A priori, it is unclear whether this condition is even satisfied in the sink state, in which *N* exhibits fluctuations that are large (relative to the mean) and characterised by significant skew towards small sizes. Figure S1b suggests that it may nevertheless be reasonable to approximate the average load in the sink state by the equilibrium expectation, at least for large values of *r*_0_*K*.

Accordingly, we note that the accuracy of the semi-deterministic prediction for the critical migration threshold, *m*_*c,*1_, at which the sink state vanishes, improves with increasing *r*_0_*K*. This is illustrated in the inset of fig. S1a, which shows *P*_*sink*_, the probability that the population is in the sink state (as observed in simulations) against *m*_0_. The semi-deterministic prediction for *m*_*c,*1_ (vertical dashed lines) approaches the corresponding threshold in simulations (where *P*_*sink*_ goes to 0) as *r*_0_*K* increases.

#### Semi-deterministic prediction for equilibrium population size in the absence of migration

In the absence of migration (i.e., with *M*_0_=0), there is always an equilibrium at *N*=0. From eq. (3) of the main text, it follows that this equilibrium is stable if 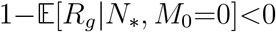. To compute the expected load in the *N*→0 limit, note that the expected frequency and the expected heterozygosity are respectively 1/2 and 0 in this limit, so that 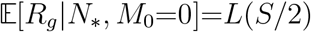. Thus, the extinction equilibrium is stable if *L*(*S/*2)>1.

There may exist a second equilibrium at 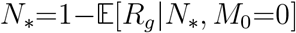; the population size *N*_*_ is positive (i.e., the population not extinct) only if 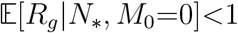 for some *N*_*_>0, i.e., if the equilibrium genetic load is lower than the baseline growth rate.

#### Semi-deterministic prediction for *m*_*c,*1_ in the *r*_0_*K*→∞ limit

In order to obtain the semi-deterministic prediction for the population size, genetic load and expected allele frequencies at one or more equilibria, we numerically solve for the {*N*_*_} satisfying:

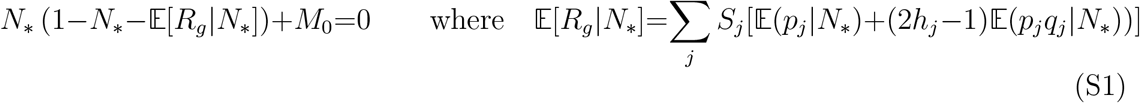

where the expectations are obtained by integrating over the equilibrium allele frequency distributions (eq. (2) of the main text). The equation above is the same as eq. (3) of the main text. An equilibrium *N*_*_ is a stable equilibrium if the function 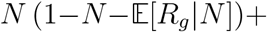 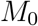 has a negative derivative with respect to *N* at *N*=*N*_*_.

In the limit *r*_0_*K*→∞, *L*→∞, *s*→0, *u*→0, with *m*_0_, *Ks*, *Ku* and 2*LU*=2*L*(*u/r*_0_) constant, the demographic contributions of migration, represented by the *M*_0_=*m*_0_*/r*_0_*K* term in the above equation, can be neglected, yielding 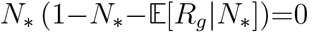. This function (which must be zero at equilibrium) is plotted as a function of population size in fig. S2a for three different levels of migration (which influence the function via the term 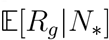). There is always an equilibrium (*N*=0) at extinction; this equilibrium is stable if the curve is downward sloping at *N*=0, i.e., if 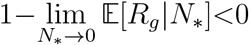. In addition, there may be two other equilibria— one stable and the other unstable— satisfying 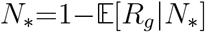. For low 2*LU* and/or*h* not too low, an increase in migration causes the sink equilibrium to become unstable, so that above a critical migration threshold *m*_*c,*1_, only a single, stable ‘large-population’ equilibrium exists.

**Figure S2:**
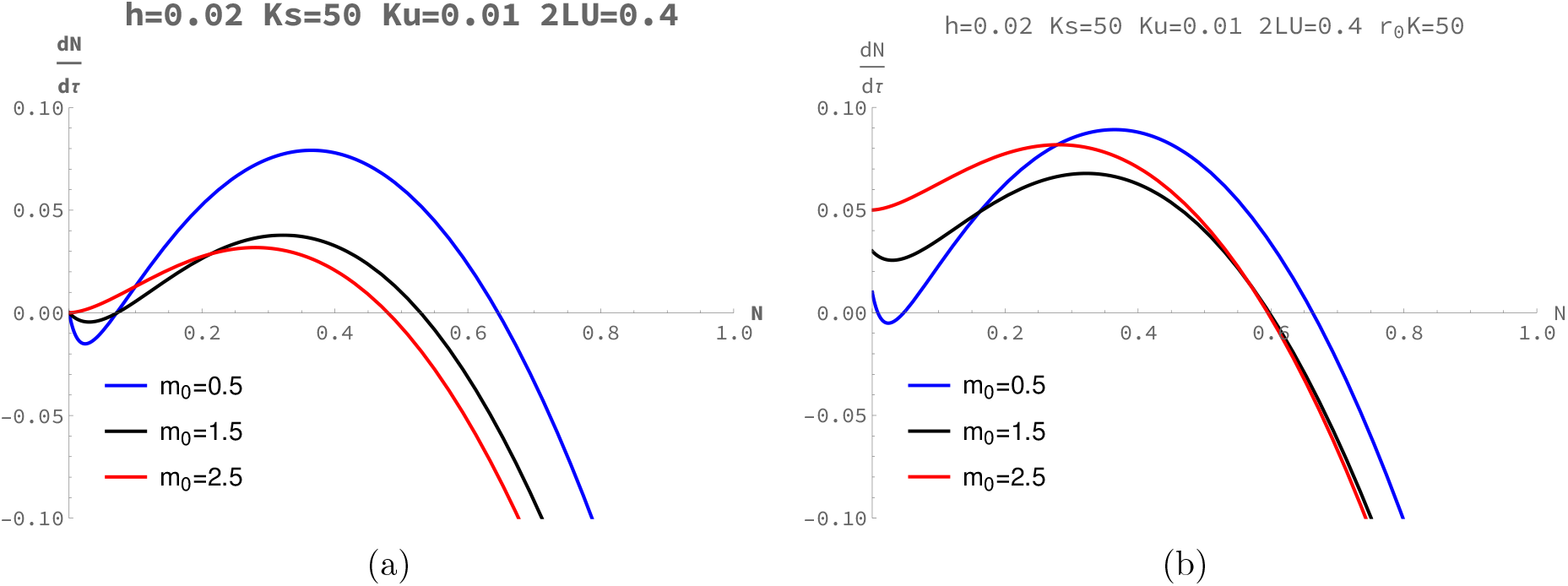
(A)-(B) Rate of change of population size *N* under the semi-deterministic approximation (eq. (S1)) as a function of *N*(A) in the *r*_0_*K*→∞ limit (where the demographic effects of migration, represented by the *M*_0_ term, can be neglected) (B) for finite *r*_0_*K*(i.e., including the *M*_0_ term). Equilibria correspond to points at which the curves cross the horizontal (zero growth rate axis); stable equilibria are those for which the curves have a negative slope at the point of zero crossing. Parameter values (*Ks*=50, *Ku*=0.01, 2*LU*=0.4 and *h*=0.02 in A and B; *r*_0_*K*=50 in B) correspond to a regime where increasing migration causes the extinction fixed point to become unstable (in the *r*_0_*K*→∞ limit; fig. A) or the sink fixed point to vanish (for finite *r*_0_*K*; fig B).

In the limit *r*_0_*K*→∞, the change in the stability properties of the sink equilibrium occurs at the migration rate for which 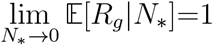. Since 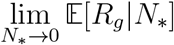, the expected load in the limit *N*_*_→0, depends only on the neutral allele frequency and neutral heterozygosity under migration-drift balance, we can obtain an explicit expression that depends on *LS*, *h*, *p*^(*m*)^ and *m*_0_. Finally, setting 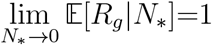 and solving for the number of migrants per generation yields:

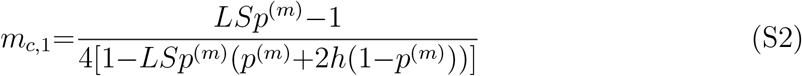

which is eq. (4) of the main text.

In order to illustrate the effects of the demographic term on equilibria and their stability properties, we also plot 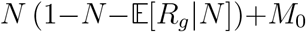 (eq. (S1) including the *M*_0_ term) as a function of *N* (fig. S2b). At low migration levels, there is a sink equilibrium (with *N*_*_ close to but not equal to zero) and a large-population equilibrium. Increasing migration now causes the sink equilibrium to vanish, rather than rendering it unstable. The critical threshold *m*_*c,*1_ in this case is significantly lower than the *r*_0_*K*→∞ prediction above (see also fig. 3B of the main text), highlighting how, migration can influence population outcomes via both genetic and demographic effects.

### C. Comparison of individual-based simulations with simulations under LE and IE

Here we compare results from the two simulation approaches by looking at the effect of migration on the mean population size and mean genetic load at equilibrium (fig. S3a – S3c) as well as on the stochastic distribution of population size (fig. S3d – S3f) on the island under different genetic architectures i.e. for weakly deleterious alleles (*Ks<Ks*; left column), mildly or moderately deleterious alleles (*Ks*≳*Ks*_*c*_; middle column) and strongly deleterious alleles (*Ks*≫*Ks*_*c*_; right column).

**Figure S3:**
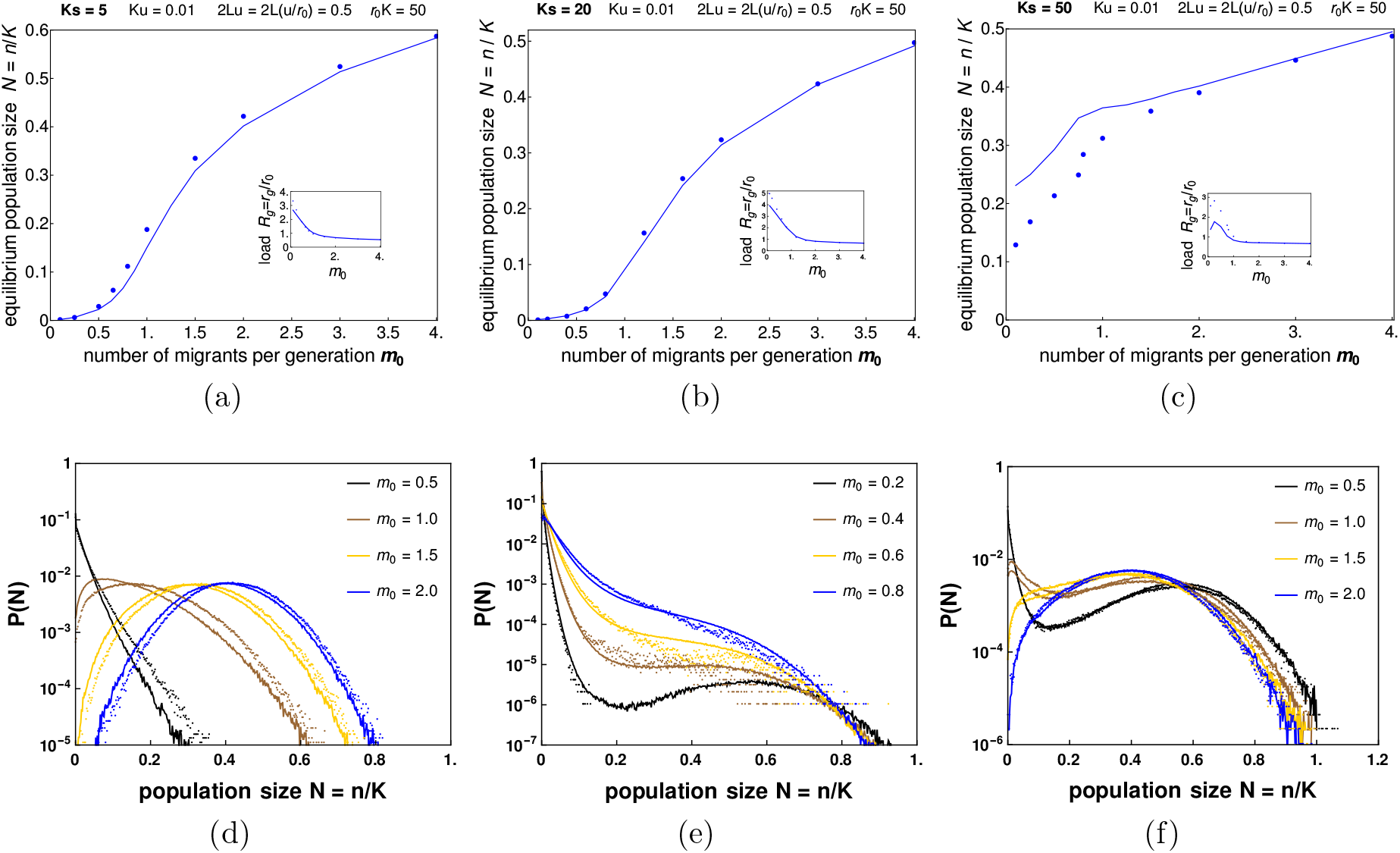
(A)-(C) Mean population size (main figure) and mean genetic load (inset) at equilibrium plotted against *m*_0_ (the number of migrants per generation) for weakly deleterious (*Ks<Ks*_*c*_; left column), mildly deleterious (*Ks*≳*Ks*_*c*_; middle column) and strongly deleterious (*Ks≫Ks*_*c*_; right column) nearly recessive alleles. Solid lines represent result obtained from allele frequency simulations (assuming LE and zero inbreeding) while dots represent result from individual-based simulations. (D)-(F) Equilibrium probability distribution of scaled population sizes *N*=*n/K* for various values of *m*_0_, as obtained from simulations assuming LE and IE (solid lines) as well as from individual based simulations (dots) under the three parameter regimes.

#### Weakly deleterious nearly recessive alleles (left column of fig. S3)

For *Ks<Ks*_*c*_, the population approaches extinction for low migration. However, increasing migration causes an increase in the size of the island population (main plot of fig. S3a) as well as a corresponding decrease in the genetic load (inset). This is true for both types of simulations although the individual based simulation (dots) produces slightly higher population numbers with increasing*m*_0_. Looking at the stochastic distribution of population size (fig. S3d), populations exhibit a single stable equilibrium for all migration rates and shift to the right towards increasing *N* (following a Gaussian distribution centered around the stable equilibrium) as *m*_0_ increases. As noticed in fig. S3a, there exists a slight difference between both simulation approaches, namely that the individual based simulation produces slightly higher numbers (see dotted plots in fig. S3d). This suggests that LD can cause a somewhat higher increase in population size by its purging effect on sets of locally maladapted alleles.

#### Moderately deleterious nearly recessive alleles (middle column of fig. S3)

Just as with weakly deleterious alleles, an increase in migration increases the population size (main plot of fig. S3b) and reduces the genetic load (inset) on the island, although we observe a higher load for low values of migration in the individual based simulation. Looking at fig. S3e, the individual based simulation breaks down for very low migration rates as it requires extremely long simulation runs (both time and memory consuming) to observe the peak at the large population state equilibrium (see LE simulations - solid lines).

#### Strongly deleterious nearly recessive alleles (right column of fig. S3)

For *Ks*≫*Ks*_*c*_, we observe an initial sharp increase in population size with *m*_0_. Beyond *m*_0_=1, the population size increases further but with a less steep slope. Similarly, for very low *m*_0_, the genetic load in the population increases initially and then falls sharply with increasing *m*_0_ after which it approaches a steady value. Interestingly, for low migration rates, individual based simulations slightly diminish the population size and increase the load (see blue dots) suggesting a negative effect of LD on strongly deleterious alleles when migration is low. From fig. S3f, we observe a bimodal distribution of population size (with one peak close to extinction and the other close to the deterministic equilibrium) when migration is rare. However, with increasing migration, there is a shift from the bimodal to a unimodal population size distribution.

### D. Evolutionary outcomes with a distribution of fitness effects

In the main text, we analyse scenarios where all deleterious variants have equal selective effects and dominance coefficients, and illustrate how migration may have qualitatively different effects, depending on the magnitude of the homozygous selection coefficient, dominance coefficient and the total mutation rate. Here, we ask: what is the effect of migration on population outcomes when the genetic load is due to loci with a distribution of fitness effects? For the purposes of illustration, we consider a somewhat artificial distribution where a fraction *α* of loci are subject to deleterious mutations that are nearly recessive (*h*_*R*_=0.02) with scaled selection coefficient *Ks*_*R*_, and the remaining fraction 1−*α* are additive (*h*_*A*_=0.5) with scaled selective coefficient *Ks*_*A*_. Thus, the mutation targets for the two types of deleterious variants are 2*LαU* and 2*L*(1−*α*)*U* respectively.

If recessive variants are weakly deleterious (*Ks*_*R*_ less than the corresponding critical threshold *Ks*_*c*_), then their contribution to genetic load will decrease with increasing migration. Since migration always reduces additive load (though only marginally for *Ks*_*A*_≫1), the net effect of increasing migration in this case is to reduce load and increase the equilibrium population size (results not shown). We thus focus on recessive alleles with moderate or strongly deleterious effects (*Ks*_*R*_>*Ks*_*c*_), where an increase in recessive load with increasing migration can potentially counteract a reduction in additive load. Figures S4a-S4c show semi-deterministic predictions for equilibrium population sizes versus *m*_0_ for various combinations of *Ks*_*A*_ and *Ks*_*R*_ for *α*=0.1 (predominantly additive load), 0.5 (nearly equal contributions of additive and recessive alleles to load) and 0.9 (load primarily due to recessive alleles). Where two stable equilibria exist, these are shown by solid and dashed lines (of the same color).

**Figure S4:**
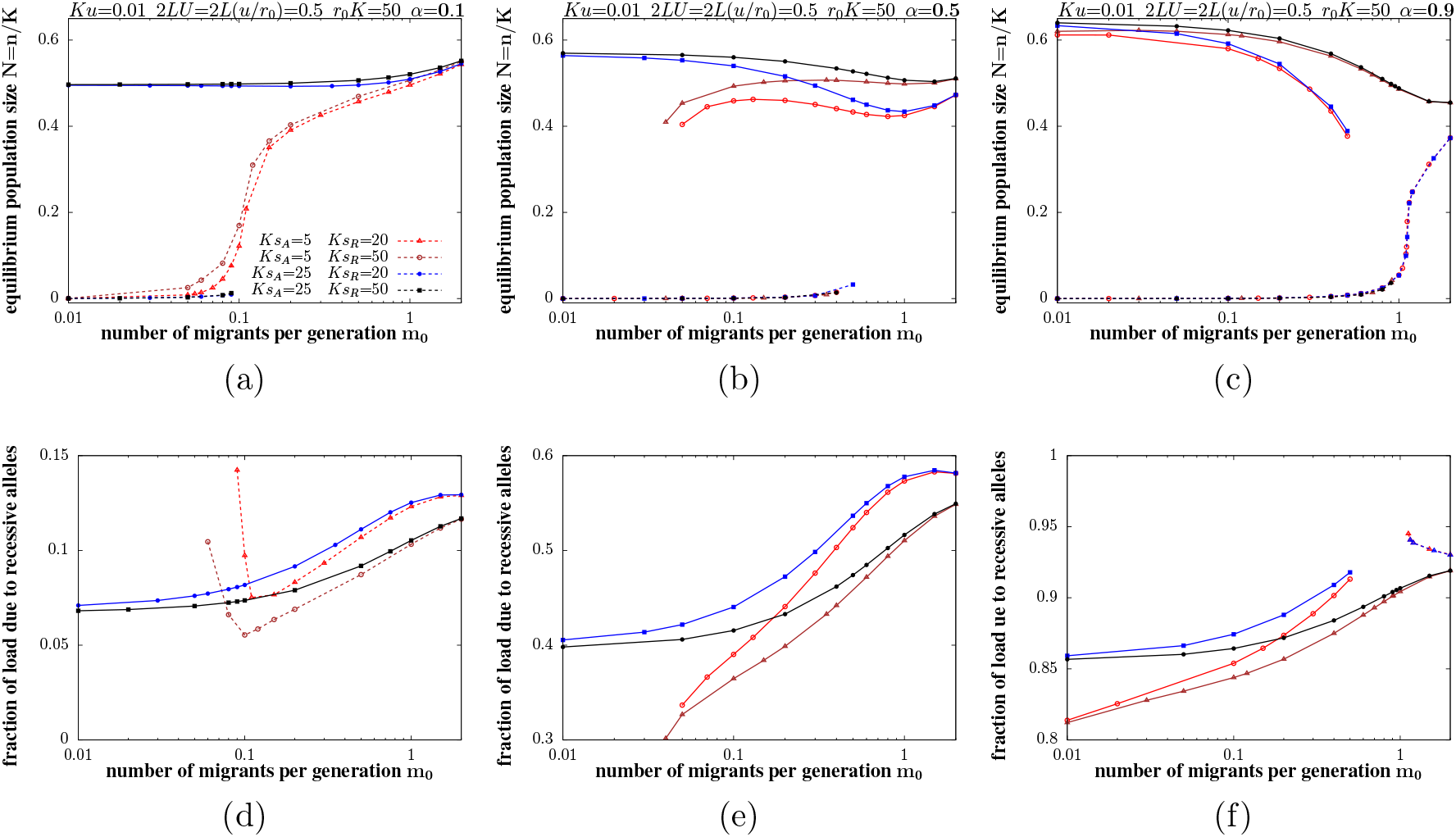
(A)-(C) Semi-deterministic predictions for population size(s) at equilibrium vs. *m*_0_, the number of migrants per generation. Different colors correspond to different values of *Ks*_*A*_ and *Ks*_*R*_, the (scaled) homozygous selection coefficients for the additive (*h*=0.5) and nearly recessive (*h*=0.02) alleles (see legend of fig. A). We depict cases where a fraction *α* of alleles have nearly recessive effects and the remaining fraction 1−*α* additive effects, for *α*=0.1 (left), *α*=0.5 (middle) and *α*=0.9 (right). Where two alternative equilibria exist, these are shown by solid and dashed lines (of the same color). (D)-(F) Semi-deterministic predictions for the fraction of the total load that is due to recessive alleles vs. *m*_0_, for parameter combinations where the total load is less then *r*_0_, i.e., where the population is not a sink. Where two equilibria with load less than *r*_0_ exist, the fractions corresponding to each are depicted by solid and dashed lines. All figures show results for: *Ku*=0.01, 2*LU*=0.5, *r*_0_*K*=50.

We observe a genetic Allee effect (characterised by the co-existence of alternative ‘sink’ and ‘large-population’ equilibria) at low migration rates for all parameter combinations, except when load is primarily due to weakly deleterious additive alleles (red and brown curves in fig. S4a). Increasing migration tends to destabilize the sink state in all cases, except where load is largely due to recessive alleles with moderately deleterious effects (red and blue curves in fig. S4c): in this extreme limit, it is the large-population state that vanishes at high migration levels (see also fig. 2C, main text). The critical migration threshold at which the sink state vanishes (and Allee effects no longer occur) is higher when the contribution of recessive alleles to total load is higher (*α* larger) and homozygous effects associated with recessive alleles more moderate. The population size associated with the large-population equilibrium increases marginally with increasing migration for small *α*(as expected when load is predominantly additive), decreases with increasing migration for large *α*(as expected when load is predominantly recessive), and exhibits a weak, non-monotonic dependence on *m*_0_ for intermediate *α*(reflecting the opposing effects of migration on additive and recessive alleles).

We also track how the relative contributions of additive and recessive alleles to total load change with increasing migration. Figures S4d-S4f shows the fraction of total load that is due to recessive alleles vs. *m*_0_, for parameter combinations where the population is not a sink, i.e., where the total load is less than the baseline growth rate*r*_0_. This fraction increases with increasing *m*_0_ in all cases (except when the population is close to a sink state, i.e., *R*_*g*_~1), with the most significant increase occurring for*α*=0.5, and where additive alleles have weak effects and recessive alleles moderate effects. Under these conditions, smaller isolated populations (with low migration) can more efficiently purge recessive load, but maintain higher levels of additive load. An increase in migration tends to decrease the additive load and increase the recessive load by pushing the frequencies of deleterious additive and recessive alleles closer to the mainland values (that are respectively lower and higher than those on the island).

Note that in this case, the total load and population size do not change significantly with migration (because of the opposing effects of migration on the two components of load). The change in the relative contributions of different kinds of alleles may nevertheless have significant consequences, by making populations with higher levels of migration (and accordingly, a greater number of segregating recessive alleles) more vulnerable to future extinction, e.g., in the event of a bottleneck.

## References

[1] Lande R. 1994. Risk of population extinction from fixation of new deleterious mutations. Evolution, vol. 48, no. 5, Oct. 1994, p. 1460. DOI.org (Crossref), doi:10.2307/2410240.

[2] Lynch, M., Conery, J., Bürger, R. (1995), Mutational meltdowns in sexual populations. Evolution, vol. 49, no. 6, Dec. 1995, p. 1067. DOI.org (Crossref), doi:10.2307/2410432.

[3] Frankham, R. (2005) Genetics and extinction. Biological Conservation, vol. 126, no. 2, Nov. 2005, pp. 131–40. DOI.org (Crossref), doi:10.1016/j.biocon.2005.05.002.

[4] O’Grady, J.J., Brook, B.W., Reed, D.H., Ballou, J.D., Tonkyn, D.W., and Frankham, R. (2006) Realistic levels of inbreeding depression strongly affect extinction risk in wild populations. Biological Conservation, vol. 133, no. 1, Nov. 2006, pp. 42–51. DOI.org (Crossref), doi:10.1016/j.biocon.2006.05.016.

[5] Higgins, K., Lynch, M. (2001) Metapopulation extinction caused by mutation accumulation. Proceedings of the National Academy of Sciences of the USA, vol. 98, no. 5, Feb. 2001, pp. 2928–33. DOI.org (Crossref), doi:10.1073/pnas.031358898.

[6] Agrawal, A.F., and Whitlock, M.C. (2011). Inferences about the distribution of dominance drawn from yeast gene knockout data. Genetics, vol. 187, no. 2, Feb. 2011, pp. 553–66. DOI.org (Crossref), doi:10.1534/genetics.110.124560.

[7] Huber, Christian D., Durvasula, A., Hancock, A.M., Lohmueller, K.E. (2018). “Gene Expression Drives the Evolution of Dominance.” Nature Communications, vol. 9, no. 1, Dec. 2018, pp. 2750. DOI.org (Crossref), doi:10.1038/s41467-018-05281-7.

[8] Lande, R. (1993). Risks of Population Extinction from Demographic and Environmental Stochasticity and Random Catastrophes. The American Naturalist, vol. 142, no. 6, Dec. 1993, pp. 911–27. DOI.org (Crossref), doi:10.1086/285580.

[9] Lande, R. (1988). Genetics and Demography in Biological Conservation. Science, vol. 241, no. 4872, Sept. 1988, pp. 1455–60. DOI.org (Crossref), doi:10.1126/science.3420403.

[10] Ovaskainen, O., and Hanski, I. Extinction Threshold in Metapopulation Models. Annales Zoologici Fennici, vol. 40, no. 2, 2003, pp. 81–97. DOI.org (Crossref), doi:10.1126/science.3420403.

[11] Gyllenberg, M., Hanski, I. Single-species metapopulation dynamics: A structured model. Theoretical Population Biology, vol. 42, no. 1, Aug. 1992, pp. 35–61. DOI.org (Crossref), doi:10.1016/0040-5809(92)90004-D.

[12] Lande, R., Engen, S., and Saether, B.-E. 1998. Extinction times in finite metapopulation models with stochastic local dynamics. Oikos, vol. 83, no. 2, Nov. 1998, p. 383. DOI.org (Crossref), doi:10.2307/3546853.

[13] Ronce, O., and Kirkpatrick, M. 2001. When sources become sinks: migrational meltdown in heterogeneous habitats. Evolution, vol. 55, no. 8, Aug. 2001, pp. 1520–31. DOI.org (Crossref), doi:10.1111/j.0014-3820.2001.tb00672.x.

[14] Szép, E., Sachdeva, H. and Barton, N.H. (2021), Polygenic local adaptation in metapopulations: A stochastic eco-evolutionary model. Evolution, vol. 75, no. 5, May 2021, pp. 1030–45. DOI.org (Crossref), doi:10.1111/evo.14210.

[15] Bell, G., and Gonzalez, A. 2011. Adaptation and evolutionary rescue in metapopulations experiencing environmental deterioration. Science, vol. 332, no. 6035, June 2011, pp. 1327–30. DOI.org (Crossref), doi:10.1126/science.1203105.

[16] Uecker, H., Otto, S.P., and Hermisson, J. 2014. Evolutionary rescue in structured populations. American Naturalist, vol. 183, no. 1, Jan. 2014, pp. E 17–35. DOI.org (Crossref), doi:10.1086/673914.

[17] Bridle JR, Vines TH. 2007 Limits to evolution at range margins: when and why does adaptation fail? Trends Ecol. Evol. vol. 22, no. 3, Mar. 2007, pp. 140–47. DOI.org (Crossref), doi:10.1016/j.tree.2006.11.002.

[18] Whitlock, M.C. (2002), Selection, Load and Inbreeding Depression in a Large Metapopulation. Genetics, vol. 160, no. 3, Mar. 2002, pp. 1191–202. DOI.org (Crossref), doi:10.1093/genetics/160.3.1191.

[19] Roze, D. 2015. Effects of interference between selected loci on the mutation load, inbreeding depression and heterosis. Genetics, vol. 201, no. 2, Oct. 2015, pp. 745–57. DOI.org (Crossref), doi:10.1534/genetics.115.178533.

[20] Lenormand, T., Gene flow and the limits to natural selection. Trends in Ecology and Evolution, vol. 17, no. 4, Apr. 2002, pp. 183–89. DOI.org (Crossref), doi:10.1016/S0169-5347(02)02497-7.

[21] Kawecki, T.J. 2008. Adaptation to marginal habitats. Annual Review of Ecology, Evolution, and Systematics, vol. 39, no. 1, Dec. 2008, pp. 321–42. DOI.org (Crossref), doi:10.1146/annurev.ecolsys.38.091206.095622.

[22] Barton, N.H., and Etheridge, A.M. 2018. Establishment in a new habitat by polygenic adaptation. Theor. Popul. Biol, vol. 122, July 2018, pp. 110–27. DOI.org (Crossref), doi:10.1016/j.tpb.2017.11.007.

[23] Wright, S., 1937 The distribution of gene frequencies in populations. Science 85: 504. DOI.org (Crossref), doi:10.1126/science.85.2212.504-a.

[24] Charlesworth, B., 2015 Causes of natural variation in fitness–evidence from studies of Drosohila populations. Proceedings of the National Academy of Sciences (U.S.A.), vol. 112, no. 6, Feb. 2015, pp. 1662–69. DOI.org (Crossref), doi:10.1073/pnas.1423275112.

[25] Charlesworth, B., 2018 Mutational load, inbreeding depression and heterosis in subdivided populations. Mol. Ecol. vol. 27, no. 24, Dec. 2018, pp. 4991–5003. DOI.org (Crossref), doi:10.1111/mec.14933.

[26] Morjan C.L., Rieseberg L.H., 2004 How species evolve collectively: implications of gene flow and selection for the spread of advantageous alleles. Mol. Ecol. vol. 13, no. 6, Mar. 2004, pp. 1341–56. DOI.org (Crossref), doi:10.1111/j.1365-294X.2004.02164.x.

[27] Hill, W. G., and A. Robertson. 1966. The effect of linkage on limits to artificial selection. Genetical Research 8:269294. DOI.org (Crossref), DOI: https://doi.org/10.1017/S0016672300010156

[28] Graur D., Zheng Y., Price N., Azevedo R. B. R., Zufall R. A., Elhaik E., 2013 On the Immortality of Television Sets: “Function” in the Human Genome According to the Evolution-Free Gospel of ENCODE. Genome Biol Evol 5: 578–590. DOI.org (Crossref), https://doi.org/10.1093/gbe/evt028.

[29] Böndel, K. B., Kraemer, S. A., Samuels, T. S., McClean, D., Lachapelle, J., Ness, R. W., Colegrave, N. and Keightley, P. D. (2019). Inferring the distribution of fitness effects of spontaneous mutations in Chlamydomonas reinhardtii. PLoS Biology 17: e3000192. DOI.org (Crossref), https://doi.org/10.1371/journal.pbio.3000192.

[30] Kimura, M., and Maruyama, T. (1966). The mutational load with epistatic gene interactions in fitness. Genetics, vol. 54, no. 6, Dec. 1966, pp. 1337–51. DOI.org (Crossref), doi:10.1093/genetics/54.6.1337.

[31] Kondrashov, A.S (1988). Deleterious mutations and the evolution of sexual reproduction. Nature, vol. 336, no. 6198, Dec. 1988, pp. 435–40. DOI.org (Crossref), doi:10.1038/336435a0.

[32] Courchamp F, Berec L, Gascoigne J (2008) Allee effects in ecology and conservation. New York: Oxford University Press.

[33] Caballero, A., Keightley, P.D. and Hill, W.G. 1991 Strategies for increasing fixation probabilities of recessive mutations. Genet. Res. vol. 58, no. 2, Oct. 1991, pp. 129–38. DOI.org (Crossref), doi:10.1017/S0016672300029785.

