## Supplementary material for "Genetic load and extinction in peripheral populations: the roles of migration, drift and demographic stochasticity": AlleleFrequencySimulation.f

```
      program wright_f2sher
      integer(KIND=8) L,K,k6,histories,i,j,j1
      integer(KIND=8) k5,tsteps,mig,nnew,c3,j3,j6,c1,mig2
      real(KIND=8) sel1,migrants,rmax,alpha,x,y,nav,t1,t2
      real(KIND=8) temp,temp1,temp2,theta,temp3,loglimit,fitness
      real(KIND=8) temp4,temp5,t2a,t3a,t4a,t1a,x2,mu,dom
      integer(KIND=8),allocatable, dimension(:)::pop,pdist
      real(KIND=8), allocatable, dimension(:)::pfreq,avgfreq,load
      real(KIND=8), allocatable, dimension(:):: avgsize,temp6
      integer(KIND=8) ZBQLBIN, ZBQLGEO, ZBQLPOI


      K=500
      L=1250
      dom=0.02d+0      
      rmax=0.1d+0
      sel1=5.0d+0/K
      pcont=0.0355724358845670058335853206971899563886d+0
      migrants=0.25D+0
      mu=0.01d+0/K
      
      loglimit=-log(tiny(double))

      histories=1
      tsteps=1000000
      theta=1.0D+0/histories


      allocate(avgfreq(tsteps),load(tsteps),pop(tsteps))
      allocate(pfreq(L),avgsize(tsteps),temp6(tsteps),pdist(2*K))

      do i=1,tsteps
      avgfreq(i)=0.0d+0
      load(i)=0.0d+0
      avgsize(i)=0.0d+0
      temp6(i)=0.0d+0
      enddo

      do i=1,2*K
      pdist(i)=0
      enddo

       call init_random_seed

       do k6=1,histories
       print*, k6, "aa"

       
       do j=1,L
       pfreq(j)=0.000d+0
       enddo         
       pop(1)=K


        do i=1,tsteps
        if (i>1) then
        pop(i)=pop(i-1)
        endif
 !      migration
         
         mig2=0
         temp1=1.0D+0
         temp3=exp(-migrants)
  3      call random_number(y)
         temp1=temp1*y
         if  (temp1>temp3)then
         mig2=mig2+1
         go to 3
         endif
         if (pop(i)+mig2>0)then
         do j=1,L
         pfreq(j)=pop(i)*pfreq(j)+mig2*pcont
         enddo
         pop(i)=pop(i)+mig2
         do j=1,L
         pfreq(j)=pfreq(j)/pop(i)
         enddo
         endif
   

!        mutation
         do j=1,L
         pfreq(j)=(1.0d+0-mu)*pfreq(j)+mu*(1.0d+0-pfreq(j))
         enddo

   

!       selection

        ! calculate mean fitness of each deme, new allele freq at each locus in each deme + new population size
        

         fitness=1.0d+0
         do j=1,L
         temp=1.0d+0-2.0d+0*pfreq(j)*(1.0d+0-pfreq(j))*(1.0d+0-
     c exp(-dom*sel1))-pfreq(j)*pfreq(j)*(1.0d+0-exp(-sel1))
         fitness=fitness*temp
         pfreq(j)=(pfreq(j)*(pfreq(j)*exp(-sel1)+
     c (1.0d+0-pfreq(j))*exp(-dom*sel1)))/temp
         enddo     
         ! stochastically generate new population sizes and allele frequencies


         ! new pop size
         temp=exp(rmax-(rmax/K)*pop(i))*fitness*pop(i)


         pop(i)=ZBQLPOI(temp)
!         new allele frequency
          if (pop(i)>0)then
          do j=1,L
          j3=ZBQLBIN(2*pop(i),pfreq(j))
          pfreq(j)=(j3*1.0d+0)/(2*pop(i))
          enddo
          endif

   

 ! calculate statistics
         temp4=0.0D+0
         temp5=0.0d+0

         do j=1,L
         temp4=temp4+pfreq(j)
         temp5=temp5+pfreq(j)*(pfreq(j)+dom*2.0d+0*(1.0d+0-pfreq(j)))
         enddo
         if (pop(i)>0)then
         temp6(i)=temp6(i)+1.0d+0
         avgfreq(i)=avgfreq(i)+theta*(temp4/L)
         load(i)=load(i)+theta*(temp5*sel1)
         endif
         avgsize(i)=avgsize(i)+theta*pop(i)
         if (i>50000)then
         pdist(pop(i)+1)=pdist(pop(i)+1)+1
         endif
         enddo
         enddo

        do i=1,tsteps
        temp6(i)=histories/temp6(i)
        avgfreq(i)=avgfreq(i)*temp6(i)
	load(i)=load(i)*temp6(i)
        enddo

 19   open(41,file='L1250h0.02_zeS5_zemu0.01_m0.25_ze50K500D.txt')
      do i=1,tsteps
      write(41,9) i,avgfreq(i),load(i),avgsize(i)
 9    Format(I8,1X,F15.9,1X,F15.9,1X,F15.9)
      enddo
      close(41)

 39   open(41,file='L1250h0.02_zeS5_zemu0.01_m0.25_ze50K500pdD.txt')
      do i=1,2*K
      write(41,8) i-1,pdist(i)
 8    Format(I8,1X,I8)
      enddo
      close(41)


   
      end
       
       
       
      SUBROUTINE init_random_seed()
      INTEGER :: z, m, clock
      INTEGER, DIMENSION(:), ALLOCATABLE :: seed, seedold
      
      CALL RANDOM_SEED(size = m)
      ALLOCATE(seed(m), seedold(m))
      
      CALL SYSTEM_CLOCK(COUNT=clock)
      
      seed = clock + 37 * (/ (z - 1, z = 1, m) /)
      CALL RANDOM_SEED(PUT = seed)
      CALL RANDOM_SEED(GET = seedold)
      
      DEALLOCATE(seed)
      END SUBROUTINE init_random_seed

      BLOCK DATA ZBQLBD01
*
*       Initializes seed array etc. for random number generator.
*       The values below have themselves been generated using the
*       NAG generator.
*
      COMMON /ZBQL0001/ ZBQLIX,B,C
      DOUBLE PRECISION ZBQLIX(43),B,C
      INTEGER(KIND=8) I
      DATA (ZBQLIX(I),I=1,43) /8.001441D7,5.5321801D8,
     +1.69570999D8,2.88589940D8,2.91581871D8,1.03842493D8,
     +7.9952507D7,3.81202335D8,3.11575334D8,4.02878631D8,
     +2.49757109D8,1.15192595D8,2.10629619D8,3.99952890D8,
     +4.12280521D8,1.33873288D8,7.1345525D7,2.23467704D8,
     +2.82934796D8,9.9756750D7,1.68564303D8,2.86817366D8,
     +1.14310713D8,3.47045253D8,9.3762426D7 ,1.09670477D8,
     +3.20029657D8,3.26369301D8,9.441177D6,3.53244738D8,
     +2.44771580D8,1.59804337D8,2.07319904D8,3.37342907D8,
     +3.75423178D8,7.0893571D7 ,4.26059785D8,3.95854390D8,
     +2.0081010D7,5.9250059D7,1.62176640D8,3.20429173D8,
     +2.63576576D8/
      DATA B / 4.294967291D9 /
      DATA C / 0.0D0 /
      END
******************************************************************
******************************************************************
******************************************************************
      SUBROUTINE ZBQLINI(SEED)
******************************************************************
*       To initialize the random number generator - either
*       repeatably or nonrepeatably. Need double precision
*       variables because integer storage can't handle the
*       numbers involved
******************************************************************
*	ARGUMENTS
*	=========
*	SEED	(integer, input). User-input number which generates
*		elements of the array ZBQLIX, which is subsequently used 
*		in the random number generation algorithm. If SEED=0,
*		the array is seeded using the system clock if the 
*		FORTRAN implementation allows it.
******************************************************************
*	PARAMETERS
*	==========
*	LFLNO	(integer). Number of lowest file handle to try when
*		opening a temporary file to copy the system clock into.
*		Default is 80 to keep out of the way of any existing
*		open files (although the program keeps searching till
*		it finds an available handle). If this causes problems,
*               (which will only happen if handles 80 through 99 are 
*               already in use), decrease the default value.
******************************************************************
      INTEGER(KIND=8) LFLNO
      PARAMETER (LFLNO=80)
******************************************************************
*	VARIABLES
*	=========
*	SEED	See above
*	ZBQLIX	Seed array for the random number generator. Defined
*		in ZBQLBD01
*	B,C	Used in congruential initialisation of ZBQLIX
*	SS,MM,}	System clock secs, mins, hours and days
*	HH,DD }
*	FILNO	File handle used for temporary file
*	INIT	Indicates whether generator has already been initialised
*
      INTEGER(KIND=8) SEED,SS,MM,HH,DD,FILNO,I
      INTEGER(KIND=8) INIT
      DOUBLE PRECISION ZBQLIX(43),B,C
      DOUBLE PRECISION TMPVAR1,DSS,DMM,DHH,DDD

      COMMON /ZBQL0001/ ZBQLIX,B,C
      SAVE INIT

*
*	Ensure we don't call this more than once in a program
*
      IF (INIT.GE.1) THEN
       IF(INIT.EQ.1) THEN
        WRITE(*,1)
        INIT = 2
       ENDIF
       RETURN
      ELSE
       INIT = 1
      ENDIF
*
*       If SEED = 0, cat the contents of the clock into a file
*       and transform to obtain ZQBLIX(1), then use a congr.
*       algorithm to set remaining elements. Otherwise take
*       specified value of SEED.
*
*>>>>>>>>>>>>>>>>>>>>>>>>>>>>>>>>>>>>>>>>>>>>>>>>>>>>>>>>>>>>>>>
*>>>>>>>	NB FOR SYSTEMS WHICH DO NOT SUPPORT THE  >>>>>>>
*>>>>>>>	(NON-STANDARD) 'CALL SYSTEM' COMMAND,    >>>>>>>
*>>>>>>>	THIS WILL NOT WORK, AND THE FIRST CLAUSE >>>>>>>
*>>>>>>>	OF THE FOLLOWING IF BLOCK SHOULD BE	 >>>>>>>
*>>>>>>>	COMMENTED OUT.				 >>>>>>>
*>>>>>>>>>>>>>>>>>>>>>>>>>>>>>>>>>>>>>>>>>>>>>>>>>>>>>>>>>>>>>>>
      IF (SEED.EQ.0) THEN
*>>>>>>>>>>>>>>>>>>>>>>>>>>>>>>>>>>>>>>>>>>>>>>>>>>>>>>>>>>>>>>>
*>>>>>>>	COMMENT OUT FROM HERE IF YOU DON'T HAVE  >>>>>>>
*>>>>>>>	'CALL SYSTEM' CAPABILITY ...		 >>>>>>>
*>>>>>>>>>>>>>>>>>>>>>>>>>>>>>>>>>>>>>>>>>>>>>>>>>>>>>>>>>>>>>>>
       CALL SYSTEM(' date +%S%M%H%j > zbql1234.tmp')
*
*       Try all file numbers for LFLNO to 999 
*
       FILNO = LFLNO
 10    OPEN(FILNO,FILE='zbql1234.tmp',ERR=11)
       GOTO 12
 11    FILNO = FILNO + 1
       IF (FILNO.GT.999) THEN
        WRITE(*,2)
        RETURN
       ENDIF
       GOTO 10
 12    READ(FILNO,'(3(I2),I3)') SS,MM,HH,DD
       CLOSE(FILNO)
       CALL SYSTEM('rm zbql1234.tmp')
       DSS = DINT((DBLE(SS)/6.0D1) * B)
       DMM = DINT((DBLE(MM)/6.0D1) * B)
       DHH = DINT((DBLE(HH)/2.4D1) * B)
       DDD = DINT((DBLE(DD)/3.65D2) * B)
       TMPVAR1 = DMOD(DSS+DMM+DHH+DDD,B)
*<<<<<<<<<<<<<<<<<<<<<<<<<<<<<<<<<<<<<<<<<<<<<<<<<<<<<<<<<<<<<<<<
*<<<<<<<<	... TO HERE (END OF COMMENTING OUT FOR 	  <<<<<<<
*<<<<<<<<	USERS WITHOUT 'CALL SYSTEM' CAPABILITY	  <<<<<<<
*<<<<<<<<<<<<<<<<<<<<<<<<<<<<<<<<<<<<<<<<<<<<<<<<<<<<<<<<<<<<<<<<
      ELSE
       TMPVAR1 = DMOD(DBLE(SEED),B)
      ENDIF
      ZBQLIX(1) = TMPVAR1
      DO 100 I = 2,43
       TMPVAR1 = ZBQLIX(I-1)*3.0269D4
       TMPVAR1 = DMOD(TMPVAR1,B)       
       ZBQLIX(I) = TMPVAR1
 100  CONTINUE

 1    FORMAT(//5X,'****WARNING**** You have called routine ZBQLINI ',
     +'more than',/5X,'once. I''m ignoring any subsequent calls.',//)
 2    FORMAT(//5X,'**** ERROR **** In routine ZBQLINI, I couldn''t',
     +' find an',/5X,
     +'available file number. To rectify the problem, decrease the ',
     +'value of',/5X,
     +'the parameter LFLNO at the start of this routine (in file ',
     +'randgen.f)',/5X,
     +'and recompile. Any number less than 100 should work.')
      END
******************************************************************
      FUNCTION ZBQLU01(DUMMY)
*
*       Returns a uniform random number between 0 & 1, using
*       a Marsaglia-Zaman type subtract-with-borrow generator.
*       Uses double precision, rather than integer, arithmetic 
*       throughout because MZ's integer constants overflow
*       32-bit integer storage (which goes from -2^31 to 2^31).
*       Ideally, we would explicitly truncate all integer 
*       quantities at each stage to ensure that the double
*       precision representations do not accumulate approximation
*       error; however, on some machines the use of DNINT to
*       accomplish this is *seriously* slow (run-time increased
*       by a factor of about 3). This double precision version 
*       has been tested against an integer implementation that
*       uses long integers (non-standard and, again, slow) -
*       the output was identical up to the 16th decimal place
*       after 10^10 calls, so we're probably OK ...
*
      DOUBLE PRECISION ZBQLU01,DUMMY,B,C,ZBQLIX(43),X,B2,BINV
      INTEGER(KIND=8) CURPOS,ID22,ID43

      COMMON /ZBQL0001/ ZBQLIX,B,C
      SAVE /ZBQL0001/
      SAVE CURPOS,ID22,ID43
      DATA CURPOS,ID22,ID43 /1,22,43/

      B2 = B
      BINV = 1.0D0/B
 5    X = ZBQLIX(ID22) - ZBQLIX(ID43) - C
      IF (X.LT.0.0D0) THEN
       X = X + B
       C = 1.0D0
      ELSE
       C = 0.0D0
      ENDIF
      ZBQLIX(ID43) = X
*
*     Update array pointers. Do explicit check for bounds of each to
*     avoid expense of modular arithmetic. If one of them is 0 the others
*     won't be
*
      CURPOS = CURPOS - 1
      ID22 = ID22 - 1
      ID43 = ID43 - 1
      IF (CURPOS.EQ.0) THEN
       CURPOS=43
      ELSEIF (ID22.EQ.0) THEN
       ID22 = 43
      ELSEIF (ID43.EQ.0) THEN
       ID43 = 43
      ENDIF
*
*     The integer arithmetic there can yield X=0, which can cause 
*     problems in subsequent routines (e.g. ZBQLEXP). The problem
*     is simply that X is discrete whereas U is supposed to 
*     be continuous - hence if X is 0, go back and generate another
*     X and return X/B^2 (etc.), which will be uniform on (0,1/B). 
*
      IF (X.LT.BINV) THEN
       B2 = B2*B
       GOTO 5
      ENDIF

      ZBQLU01 = X/B2

      END
******************************************************************
      FUNCTION ZBQLUAB(A,B)
*
*       Returns a random number uniformly distributed on (A,B)
*
      DOUBLE PRECISION A,B,ZBQLU01,ZBQLUAB
      
*
*       Even if A > B, this will work as B-A will then be -ve
*
      IF (A.NE.B) THEN
       ZBQLUAB = A + ( (B-A)*ZBQLU01(0.0D0) )
      ELSE
       ZBQLUAB = A
       WRITE(*,1)
      ENDIF
 1    FORMAT(/5X,'****WARNING**** (function ZBQLUAB) Upper and ',
     +'lower limits on uniform',/5X,'distribution are identical',/)
      END
******************************************************************
      FUNCTION ZBQLEXP(MU)
*
*       Returns a random number exponentially distributed with
*       mean MU
*
      DOUBLE PRECISION MU,ZBQLEXP,ZBQLU01

      ZBQLEXP = 0.0D0

      IF (MU.LT.0.0D0) THEN
       WRITE(*,1)
       RETURN
      ENDIF

      ZBQLEXP = -DLOG(ZBQLU01(0.0D0))*MU

 1    FORMAT(/5X,'****ERROR**** Illegal parameter value in ',
     +' ZBQLEXP',/)

      END
******************************************************************
      FUNCTION ZBQLNOR(MU,SIGMA)
*
*       Returns a random number Normally distributed with mean
*       MU and standard deviation |SIGMA|, using the Box-Muller
*       algorithm
*
      DOUBLE PRECISION THETA,R,ZBQLNOR,ZBQLU01,PI,MU,SIGMA
      DOUBLE PRECISION SPARE
      INTEGER(KIND=8) STATUS
      SAVE STATUS,SPARE,PI
      DATA STATUS /-1/

      IF (STATUS.EQ.-1) PI = 4.0D0*DATAN(1.0D0)

      IF (STATUS.LE.0) THEN
       THETA = 2.0D0*PI*ZBQLU01(0.0D0)
       R = DSQRT( -2.0D0*DLOG(ZBQLU01(0.0D0)) )
       ZBQLNOR = (R*DCOS(THETA))
       SPARE = (R*DSIN(THETA))
       STATUS = 1
      ELSE
       ZBQLNOR = SPARE
       STATUS = 0
      ENDIF
      
      ZBQLNOR = MU + (SIGMA*ZBQLNOR)

      END
******************************************************************
      FUNCTION ZBQLBIN(N,P)
*
*       Returns a random number binomially distributed (N,P)
*
      DOUBLE PRECISION P,ZBQLBET1
      DOUBLE PRECISION PP,PPP,G,Y,TINY

      INTEGER(KIND=8) N,ZBQLBIN,ZBQLGEO,IZ,NN

      TINY = 1.0D-8
      ZBQLBIN = 0

      IF (.NOT.( (P.GE.0.0D0).AND.(P.LE.1.0D0) )) THEN
       WRITE(*,1)
       RETURN
      ELSEIF(N.LE.0) THEN
       WRITE(*,1)
       RETURN
      ENDIF
*
*	First step: if NP > 10, say, things will be expensive, and 
*	we can get into the right ballpark by guessing a value for
*	ZBQLBIN (IZ, say), and simulating Y from a Beta distribution 
*	with parameters IZ and NN-IZ+1 (NN starts off equal to N).
*	If Y is less than PP (which starts off as P) then the IZth order 
*	statistic from NN U(0,1) variates is less than P, and we know 
*	that there are at least IZ successes. In this case we focus on 
*	the remaining (NN-IZ) order statistics and count how many are
*	less than PP, which is binomial (NN-IZ,(PP-Y)/(1-Y)). 
*	Otherwise, if Y is greater than PP there must be less 
*	than IZ successes, so we can count the number of order statistics
*	under PP, which is binomial (IZ-1,P/Y). When we've got NN*PP
*	small enough, we go to the next stage of the algorithm and 
*	generate the final bits directly.
*
      NN = N
      PP = P
 10   IZ = INT(DBLE(NN)*PP) + 1
      IF ( (IZ.GT.10).AND.(IZ.LT.NN-10) ) THEN
       Y = ZBQLBET1(DBLE(IZ),DBLE(NN-IZ+1))
       IF (Y.LT.PP) THEN
        ZBQLBIN = ZBQLBIN + IZ
        NN = NN - IZ
        PP = (PP-Y) / (1.0D0-Y)
       ELSE
        NN = IZ-1
        PP = PP/Y
       ENDIF
       GOTO 10
      ENDIF
*
*	PP is the probability of the binomial we're currently
*	simulating from. For the final part, we simulate either number 
*	of failures or number of success, depending which is cheaper.
*      
 20   IF (PP.GT.0.5) THEN
       PPP = 1.0D0-PP
      ELSE
       PPP = PP
      ENDIF

      G = 0
      IZ = 0
*
*     ZBQLGEO falls over for miniscule values of PPP, so ignore these
*     (tiny probability of any successes in this case, anyway)
* 
      IF (PPP.GT.TINY) THEN
 30    G = G + ZBQLGEO(PPP)
       IF (G.LE.NN) THEN
        IZ = IZ + 1
        GOTO 30
       ENDIF
      ENDIF

      IF (PP.GT.0.5) IZ = NN - IZ
      ZBQLBIN = ZBQLBIN + IZ

 1    FORMAT(/5X,'****ERROR**** Illegal parameter value in ',
     +' ZBQLBIN',/)
      END
******************************************************************
      FUNCTION ZBQLGEO(P)
*
*       Returns a random number geometrically distributed with 
*       parameter P ie. mean 1/P
* 
  
      DOUBLE PRECISION P,ZBQLU01,U,TINY
      INTEGER(KIND=8) ZBQLGEO

      TINY = 1.0D-12
      ZBQLGEO = 0
 
      IF (.NOT.( (P.GE.0.0D0).AND.(P.LE.1.0D0) )) THEN
       WRITE(*,1)
       RETURN
      ENDIF

      IF (P.GT.0.9D0) THEN
 10    ZBQLGEO = ZBQLGEO + 1 
       U = ZBQLU01(0.0D0)
       IF (U.GT.P) GOTO 10
      ELSE
       U = ZBQLU01(0.0D0)
*
*	For tiny P, 1-p will be stored inaccurately and log(1-p) may
*	be zero. In this case approximate log(1-p) by -p
*
       IF (P.GT.TINY) THEN
        ZBQLGEO = 1 + INT( DLOG(U)/DLOG(1.0D0-P) )
       ELSE
        ZBQLGEO = 1 + INT(-DLOG(U)/P)
       ENDIF
      ENDIF

 1    FORMAT(/5X,'****ERROR**** Illegal parameter value in ',
     +' ZBQLGEO',/)
      END
******************************************************************
      FUNCTION ZBQLPOI(MU)
*
*       Returns a random number Poisson distributed with mean MU
*
      
      DOUBLE PRECISION ZBQLU01,X,Y,MU,PI
      DOUBLE PRECISION ZBQLLG,ZBQLGAM,MU1,TMP1,TMP2,T
      INTEGER(KIND=8) ZBQLPOI,ZBQLBIN,K,INIT
      SAVE INIT,PI
      DATA INIT /0/

      IF (INIT.EQ.0) THEN
       PI = 4.0D0*DATAN(1.0D0)
       INIT = 1
      ENDIF

      ZBQLPOI = 0

      IF (MU.LT.0.0D0) THEN
       WRITE(*,1)
       RETURN
      ENDIF
*
*      For small MU, generate exponentials till their sum exceeds 1
*      (equivalently, uniforms till their product falls below e^-MU)
*
      IF (MU.LE.1.0D3) THEN
       MU1 = MU
*
*     For values of MU less than 1000, use order statistics - the Kth
*     event in a Poisson process of rate MU has a Gamma distribution
*     with parameters (MU,K); if it's greater than 1 we know that there 
*     are less than K events in (0,1) (and the exact number is binomial)
*     and otherwise the remaining number is another Poisson. Choose K so
*     that we'll get pretty close to 1 in the first go but are unlikely
*     to overshoot it.
*
 19    IF (MU1.GT.1.0D1) THEN        
        K = INT(MU1-DSQRT(MU1))
        Y = ZBQLGAM(DBLE(K),MU1)
        IF (Y.GT.1.0D0) THEN
         ZBQLPOI = ZBQLPOI + ZBQLBIN(K-1,(1.0D0/Y))
         RETURN
        ENDIF
        ZBQLPOI = ZBQLPOI + K
        MU1 = MU1  * (1.0D0-Y)
        GOTO 19
       ENDIF
       Y = DEXP(-MU1)
       X = 1.0D0
 20    X = X*ZBQLU01(0.0D0)
       IF (X.GT.Y) THEN
        ZBQLPOI = ZBQLPOI + 1
        GOTO 20
       ENDIF
*
*     For really huge values of MU, use rejection sampling as in 
*     Press et al (1992) - large numbers mean some accuracy may be
*     lost, but it caps the execution time.
*
      ELSE
       TMP1 = DSQRT(2.0D0*MU)
       TMP2 = ZBQLLG(MU+1.0D0)-(MU*DLOG(MU))
 30    Y = DTAN(PI*ZBQLU01(0.0D0))
       ZBQLPOI = INT(MU + (TMP1*Y))
       IF (ZBQLPOI.LT.0) GOTO 30
       X = DBLE(ZBQLPOI)
       T = (X*DLOG(MU)-ZBQLLG(X+1.0D0)) + TMP2
       IF (DABS(T).LT.1.0D2) THEN
        T = 0.9D0*(1.0D0+(Y*Y))*DEXP(T)
        IF (ZBQLU01(0.0D0).GT.T) GOTO 30
       ELSE
        T = DLOG(0.9D0*(1.0D0+(Y*Y))) + T
        IF (DLOG(ZBQLU01(0.0D0)).GT.T) GOTO 30
       ENDIF
      ENDIF 

 1    FORMAT(/5X,'****ERROR**** Illegal parameter value in ',
     +' ZBQLPOI',/)
      END
******************************************************************
      FUNCTION ZBQLGAM(G,H)
*
*       Returns a random number with a gamma distribution with mean
*       G/H and variance G/(H^2). (ie. shape parameter G & scale
*       parameter H)
*
      DOUBLE PRECISION C,D,R,ZBQLGAM,ZBQLU01,G,H,A,z1,z2,B1,B2,M
      DOUBLE PRECISION U1,U2,U,V,TEST,X
      double precision c1,c2,c3,c4,c5,w

      ZBQLGAM = 0.0D0

      IF ( (G.LE.0.0D0).OR.(H.LT.0.0D0) ) THEN
       WRITE(*,1)
       RETURN
      ENDIF

      IF (G.LT.1.0D0) THEN
889    u=ZBQLU01(0.0d0)
       v=ZBQLU01(0.0d0)
       if (u.gt.exp(1.0d0)/(g+exp(1.0d0))) goto 891
       ZBQLGAM=((g+exp(1.0d0))*u/exp(1.0d0))**(1.0d0/g)
       if (v.gt.exp(-ZBQLGAM)) then
	goto 889
       else
	goto 892
       endif
891    ZBQLGAM=-log((g+exp(1.0d0))*(1.0d0-u)/(g*exp(1.0d0)))
       if (v.gt.ZBQLGAM**(g-1.0)) goto 889
892    ZBQLGAM=ZBQLGAM/h
       RETURN
      ELSEIF (G.LT.2.0D0) THEN
       M = 0.0D0
      elseif (g.gt.10.0d0) then
       c1=g-1.0d0
       c2=(g-1.0d0/(6.0d0*g))/c1
       c3=2.0d0/c1
       c4=c3+2.0d0
       c5=1.0d0/sqrt(g)
777    u=ZBQLU01(0.0d0)
       v=ZBQLU01(0.0d0)
       if (g.gt.2.50d0) then
	u=v+c5*(1.0d0-1.860d0*u)
       endif 
       if (u.le.0.0d0.or.u.ge.1.0d0) goto 777 
       w=c2*v/u 
       if (c3*u+w+1.0d0/w.le.c4) goto 778 
       if (c3*log(u)-log(w)+w.ge.1.0d0) goto 777
778    ZBQLGAM=c1*w/h 
       return
      ELSE
       M = -(G-2.0D0) 
      ENDIF
      R = 0.50D0
      a = ((g-1.0d0)/exp(1.0d0))**((g-1.0d0)/(r+1.0d0))
      C = (R*(M+G)+1.0D0)/(2.0D0*R)
      D = M*(R+1.0D0)/R
      z1 = C-DSQRT(C*C-D)
*
*     On some systems (e.g. g77 0.5.24 on Linux 2.4.24), C-DSQRT(C*C)
*     is not exactly zero - this needs trapping if negative.
*
      IF ((Z1-M.LT.0.0D0).AND.(Z1-M.GT.-1.0D-12)) Z1 = M
      z2 = C+DSQRT(C*C-D)
      B1=(z1*(z1-M)**(R*(G-1.0D0)/(R+1.0D0)))*DEXP(-R*(z1-M)/(R+1.0D0))
      B2=(z2*(z2-M)**(R*(G-1.0D0)/(R+1.0D0)))*DEXP(-R*(z2-M)/(R+1.0D0))
50    U1=ZBQLU01(0.0D0)
      U2=ZBQLU01(0.0D0)
      U=A*U1
      V=B1+(B2-B1)*U2
      X=V/(U**R)
      IF (X.LE.M) GOTO 50
      TEST = ((X-M)**((G-1)/(R+1)))*EXP(-(X-M)/(R+1.0D0))
      IF (U.LE.TEST) THEN
       ZBQLGAM = (X-M)/H
      ELSE
       GOTO 50
      ENDIF
 1    FORMAT(/5X,'****ERROR**** Illegal parameter value in ',
     +' ZBQLGAM',/5X, '(both parameters must be positive)',/)

      END
***************************************************************
      FUNCTION ZBQLBET1(NU1,NU2)
*
*       Returns a random number, beta distributed with degrees
*       of freedom NU1 and NU2.
*
      DOUBLE PRECISION NU1,NU2,ZBQLGAM,ZBQLBET1,ZBQLU01,X1,X2

      ZBQLBET1 = 0.0D0

      IF ( (NU1.LE.0.0).OR.(NU2.LE.0.0) ) THEN
       WRITE(*,1)
       RETURN
      ENDIF
*      
*       If parameters are too small, gamma subroutine tends to return zero
*       as all the probability goes to the origin and we get rounding
*       errors, even with double precision. In this case, we use Johnk's
*       method, suitably scaled to avoid rounding errors as much as possible.
*
      
      IF ( (NU1.LT.0.9D0).AND.(NU2.LT.0.9D0) ) THEN
 10    X1 = ZBQLU01(0.0D0)
       X2 = ZBQLU01(0.0D0)
       IF ( (X1**(1.0D0/NU1))+(X2**(1.0D0/NU2)).GT.1.0D0) GOTO 10    
       X1 = (DLOG(X2)/NU2) - (DLOG(X1)/NU1)
       ZBQLBET1 = (1.0D0 + DEXP(X1))**(-1)
       IF (ZBQLBET1.GT.1.0D0) GOTO 10    
      ELSE
       X1 = ZBQLGAM(NU1,1.0D0)
       X2 = ZBQLGAM(NU2,1.0D0)
       ZBQLBET1 = X1/(X1+X2)
      ENDIF
       
      RETURN

 1    FORMAT(/5X,'****ERROR**** Illegal parameter value in ',
     +' ZBQLBET1',/5X,
     +'(both degrees of freedom must be positive)',/)

      END
***************************************************************
      FUNCTION ZBQLWEI(A,B)
*
*       Returns a random number, Weibull distributed with shape parameter
*       A and location parameter B, i.e. density is
*	f(x) = ( A/(B**A) ) * x**(A-1) * EXP( -(x/B)**A )
*
      DOUBLE PRECISION A,B,ZBQLU01,ZBQLWEI,U

      ZBQLWEI = 0.0D0

      IF ( (A.LE.0.0).OR.(B.LE.0.0) ) THEN
       WRITE(*,1)
       RETURN
      ENDIF
 
      U = ZBQLU01(0.0D0)
      ZBQLWEI = B * ( (-DLOG(U))**(1.0D0/A) )

 1    FORMAT(/5X,'****ERROR**** Illegal parameter value in ',
     +' ZBQLWEI',/5X,
     +'(both parameters must be positive)',/)
      END
***************************************************************
      FUNCTION ZBQLNB(R,P)
*
*       Returns a pseudo-random number according to a Negative
*	Binomial distribution with parameters (R,P). NB these are
*	both DOUBLE - it copes with non-integer R as well. The
*       form of the distribution is *not* the no. of trials to 
*       the Rth success - see documentation for full spec.
*
      DOUBLE PRECISION R,P,ZBQLGAM,Y
      INTEGER(KIND=8) ZBQLNB,ZBQLPOI

      ZBQLNB = 0

      IF ( (R.LE.0.0D0).OR.(P.LE.0.0D0).OR.(P.GE.1.0D0) ) THEN
       WRITE(*,1)
       RETURN
      ENDIF

      Y = ZBQLGAM(R,1.0D0)
      Y = Y*P/(1.0D0-P)
      ZBQLNB = ZBQLPOI(Y)

 1    FORMAT(/5X,'****ERROR**** Illegal parameter value in ',
     +' ZBQLNB')
      END
***************************************************************
      FUNCTION ZBQLPAR(A,B)
*
*     Returns a random number, Pareto distributed with parameters
*     A and B. The density is A*(B**A) / (B+X)**(A+1) for X > 0.
*     (this is slightly nonstandard - see documentation in 
*     randgen.txt). The algorithm is straightforward - it uses the
*     inverse CDF method.
*
      DOUBLE PRECISION A,B,ZBQLPAR,ZBQLU01,U

      ZBQLPAR = 0.0D0

      IF ( (A.LE.0.0D0).OR.(B.LE.0.0D0) ) THEN
       WRITE(*,1)
       RETURN
      ENDIF
 
      U = ZBQLU01(0.0D0)
      ZBQLPAR = B * (U**(-1.0D0/A)-1.0D0)

 1    FORMAT(/5X,'****ERROR**** Illegal parameter value in ',
     +' ZBQLPAR',/5X,
     +'(both parameters must be positive)',/)
      END
***************************************************************
      FUNCTION ZBQLLG(X)
*
*     Returns log(G(X)) where G is the Gamma function. The algorithm is
*     that given in Press et al (1992), Section 6.1, although this
*     version also allows for arguments less than 1.
*
      DOUBLE PRECISION X,Z,Z2,ZBQLLG,PI,RLN2P,C(0:6),TMP,SUM
      INTEGER(KIND=8) INIT,I
      SAVE INIT,C,RLN2P,PI
      DATA INIT /0/
      DATA (C(I),I=0,6) / 
     +              1.000000000190015D0,76.18009172947146D0,
     +              -86.50532032941677D0,24.01409824083091D0,
     +              -1.231739572450155D0,0.1208650973866179D-2,
     +              -0.5395239384953D-5/

      IF (INIT.EQ.0) THEN
        PI = 4.0D0*DATAN(1.0D0)
        RLN2P = 0.5D0*DLOG(2.0D0*PI)
        INIT = 1
      ENDIF
*
*     Compute for x > 1, then use transformation if necessary. Z is
*     our working argument.
*
      IF (X.GE.1.0D0) THEN
       Z = X
      ELSE 
       Z = 2.0D0-X
       Z2 = 1.0D0-X
      ENDIF

      IF (DABS(Z-1.0D0).LT.1.0D-12) THEN
       ZBQLLG = 0.0D0
       RETURN
      ENDIF

      TMP = Z + 4.5D0
      TMP = ( (Z-0.5D0)*DLOG(TMP) ) - TMP + RLN2P

      SUM = C(0)
      DO 50 I=1,6
       SUM = SUM + (C(I)/(Z+DBLE(I-1)))
 50   CONTINUE
      ZBQLLG = TMP + DLOG(SUM)
*
*     Transformation required if X<1
*
      IF (X.LT.1.0D0) THEN
       TMP = PI*Z2
       ZBQLLG = DLOG(TMP/DSIN(TMP)) - ZBQLLG
      ENDIF

      END
```
